# The combined action of Esrrb and Nr5a2 is essential for naïve pluripotency

**DOI:** 10.1101/2020.06.05.134999

**Authors:** Nicola Festuccia, Nick Owens, Almira Chervova, Agnès Dubois, Pablo Navarro

## Abstract

The maintenance of pluripotency in mouse embryonic stem cells (ESCs) is governed by the action of an interconnected network of transcription factors. Among them, only Oct4 and Sox2 have been shown to be strictly required for the self-renewal of ESCs and pluripotency, particularly in culture conditions where differentiation cues are chemically inhibited. Here, we report that the conjunct activity of two orphan nuclear receptors, Esrrb and Nr5a2, parallels the importance of that of Oct4 and Sox2 in naïve ESCs. By occupying a large common set of regulatory elements, these two factors control the binding of Oct4, Sox2 and Nanog to DNA. Consequently, in their absence the pluripotency network collapses and the transcriptome is substantially deregulated, leading to the differentiation of ESCs. Altogether, this work identifies orphan nuclear receptors, previously thought to be performing supportive functions, as a new set of core regulators of naïve pluripotency.

## Introduction

The uncommitted identity of mouse Embryonic Stem (ES) cells is maintained by the activity of a gene regulatory network that strictly depends on the function of Oct4 and Sox2 **(Yeo and Ng, 2013)**. Binding DNA in complex with Sox2 **(Tapia et al., 2015)**, Oct4 enables the recruitment of other transcription factors (TFs) at regulatory regions **(King and Klose, 2017)**. In accord, altering the expression of Oct4 or Sox2 results in the differentiation of ESCs. In line with their role in ESCs, Oct4 and Sox2 are essential for epiblast specification **(Avilion et al., 2003; Nichols et al., 1998)**. Alongside, a number of auxiliary factors stabilise the pluripotency network. While their individual depletion often affects the efficiency of self-renewal **(Chambers et al., 2007)** and modulates Oct4 and Sox2 binding at a subset of regulatory elements **(Heurtier et al., 2019)**, the loss of these TFs does not result in overt differentiation. For these reasons, it is generally considered that the pluripotency network is structured along two distinct modules: the core network centred on the master regulators Oct4 and Sox2, and a cohort of supportive transcription factors.

Among all auxiliary TFs, Esrrb is prominent **(Festuccia et al., 2018b)**, controlling multiple aspects of the molecular wiring of ESC identity: Esrrb is a pivotal mediator of proself-renewing signalling cues, operating downstream of the canonical WNT pathway **(Martello et al., 2012)**, and bypassing the dependence of ESCs on the LIF cytokine **(Festuccia et al., 2012)**; it acts as a major gatekeeper of pluripotency, both during early differentiation **(Festuccia et al., 2018a)** and, conversely, during reprogramming to induced pluripotency **(Adachi et al., 2018)**; it orchestrates the recruitment of the transcriptional machinery **(Bell et al., 2020; Percharde et al., 2012)** and other transcription factors **(Adachi et al., 2018)** to key regulatory elements of the pluripotency network **(Whyte et al., 2013)**; it maintains its binding activity during mitosis, directly contributing to the stability of pluripotency during cell division **(Festuccia et al., 2016)**. Given these characteristics, it is therefore not surprising that the genetic ablation of Esrrb compromises ESCs self-renewal in conventional culture conditions **(Festuccia et al., 2012)**. In contrast, the loss of Esrrb, although detrimental **(Atlasi et al., 2019)**, is tolerated in culture conditions that more stringently enforce the maintenance of the undifferentiated state **(Martello et al., 2012)** and which involve blocking the differentiation cues imparted by ERK signalling and reinforcing the activation of WNT by GSK3b inhibition **(Ying et al., 2008)**. How ESCs accommodate the concomitant disruption of the multiple aspects of Esrrb action remains unclear, representing a major gap in our knowledge of the molecular control of ESCs.

One simple hypothesis to molecularly explain how 2i/LIF can bypass the requirements for Esrrb activity is that other TFs might be performing a compensatory role. Of note, Esrrb is part of a broad family of TFs, orphan nuclear receptors, that show high sequence and structural homology, and play overlapping developmental functions **(Festuccia et al., 2018b)**. Among these, Nr5a2 interacts with Esrrb **(van den Berg et al., 2010)** and binds to nearly identical half-palindromic sequences through a related DNA binding domain presenting a common extension to the conventional zinc-finger of nuclear receptors **(Heng et al., 2010; Solomon et al., 2005)**. Further, Nr5a2 is expressed in ESCs, where it contributes to supporting, or instating, pluripotency **(Guo and Smith, 2010; Heng et al., 2010)**. To test whether Nr5a2 mitigates the consequences of loss of Esrrb function, we concomitantly ablated both TFs. Here, we report that the loss of both Esrrb and Nr5a2 leads to the complete abrogation of self-renewal and the induction of differentiation even in 2i/LIF. These effects are mediated by the full collapse of the pluripotency network: Oct4, Sox2 and Nanog binding is acutely lost at most ESC enhancers and ESC-specific gene expression is shut down. Our results identify Esrrb and Nr5a2 as a new set of essential regulators of pluripotency, whose collective importance parallels that of Oct4 and Sox2 in ESCs.

## Results

### Esrrb and Nr5a2 are co-expressed in individual ES cells and bind together at regulatory elements

Expression of auxiliary TFs, including Esrrb, is heterogeneous in ESCs cultured in serum and LIF (FCS/LIF), which induce a metastable state permissive for spontaneous differentiation **(Chambers et al., 2007)**. Notably, the loss of Esrrb in this context marks the irreversible commitment to the dismantling of pluripotency, triggering the reorganization of Oct4 binding **(Festuccia et al., 2018a)**. Therefore, we first compared the levels of expression of Esrrb and Nr5a2 in this context. Using GFP knocked-in in frame of Nr5a2 or Esrrb, linked by a self-cleaving T2a peptide (Fig S1), we observed that Esrrb presents a broad distribution of expression levels in ESCs (Fig 1A), as previously reported. Nr5a2 expression is approximately five-fold weaker, in good agreement with gene expression analysis (Fig S2A). Importantly, both genes are undetectable in a fraction of the ESC population. Next, we derived additional reporter lines in which the coding sequence for GFP and mCherry are knocked-in in frame of the Nr5a2 and Esrrb ORF, respectively (Fig S1), and found that Esrrb and Nr5a2 proteins show broadly overlapping patterns of expression, with cells negative for one TF being also low or negative for the second (Fig 1B, C). Nonetheless, possibly as a consequence of the lower expression levels, Nr5a2 down-regulation is overall more frequently observed, as confirmed by single-cell gene expression analysis (Fig 1C, S2B,C). In 2i/LIF cultures spontaneous differentiation is suppressed, and the expression of Esrrb and other auxiliary TFs is reinforced, becoming homogeneous. In line, double reporters revealed how in chemically defined medium Esrrb and Nr5a2 protein levels are elevated and their spontaneous downregulation in individual cells impeded (Fig 1B, S2B,C). These results suggest that Esrrb and Nr5a2 respond similarly to signalling cues in ESCs. Since the downregulation of Esrrb marks the commitment of ESCs to differentiate, it is possible that Nr5a2 is also relevant in this context, with the concomitant down-regulation of both TFs possibly playing a causal role in the extinction of pluripotency.

**Fig. 1.**
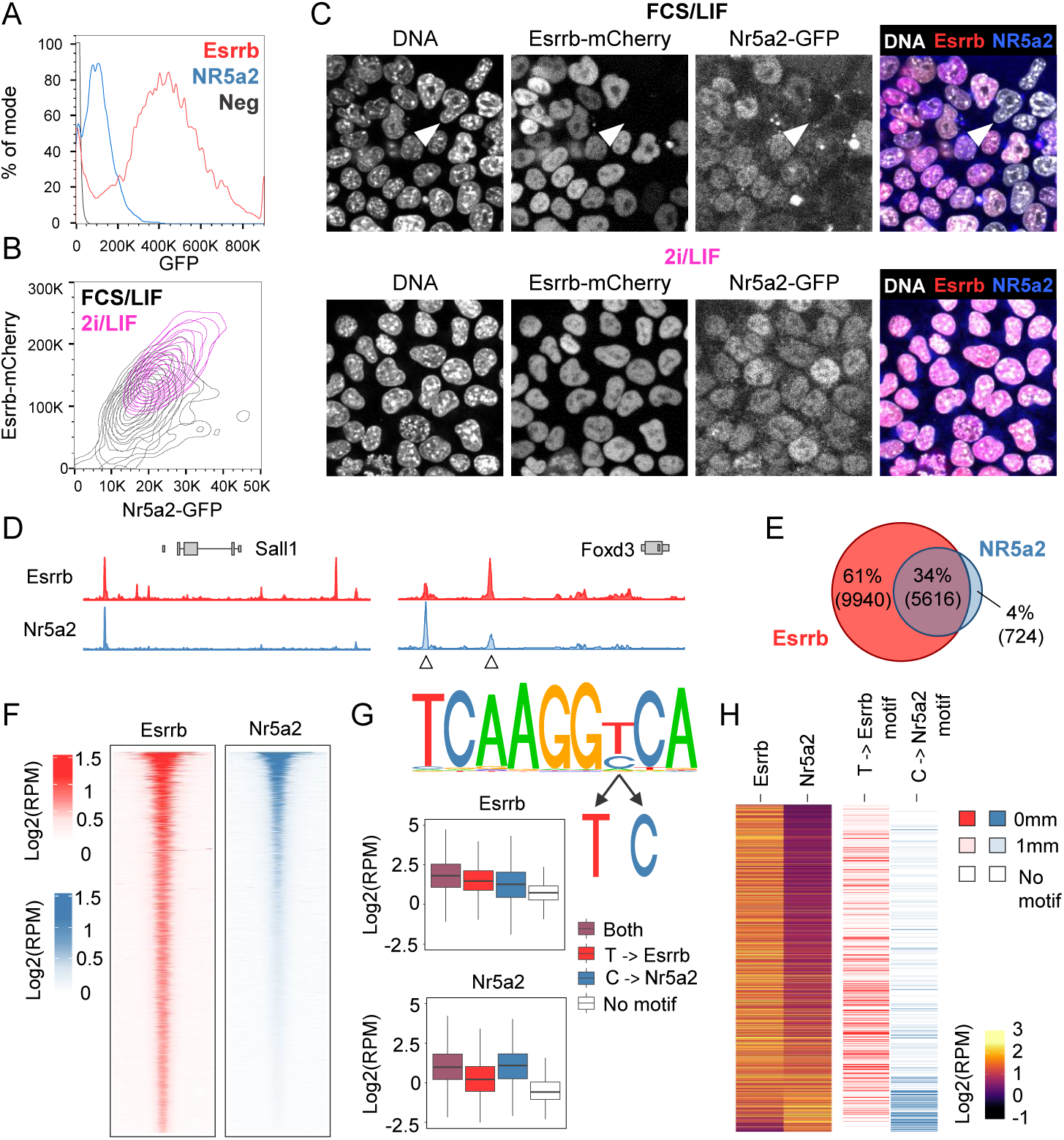
Overlapping expression and binding patterns of Esrrb and Nr5a2. **(A)** Esrrb-T2a-GFP (red) and Nr5a2-T2a-GFP (blue) fluorescence levels (percentage of mode in each dataset) determined by flow cytometry in ESC cultured in FCS/LIF. In black, background levels measured in wild-type E14Tg2a ESCs. **(B)** Nr5a2-GFP and Esrrb-mCherry levels in double knock-in ESCs cultured in FCS/LIF (black) or 2i/LIF (magenta), as determined by imaging flow cytometry (ImageStream). **(C)** Confocal microscopy images showing Esrrb-mCherry and Nr5a2-GFP expression in double knock-in ESCs cultured in FCS/LIF (top) or 2i/LIF (bottom). Note double-negative ESCs (white arrow-heads) exist only in FCS/LIF. **(D)** Profiles of Esrrb and Nr5a2 binding as determined by ChIP-seq in proximity of the Sall1 and Foxd3 genes, highlighting examples of the binding preference of each TF at common targets (arrowheads). **(E)** Venn diagram showing the intersection between regions called as peaks for Esrrb and Nr5a2 (detected in all replicates, see Methods for details). **(F)** Local enrichment heatmap comparing Esrrb (left, red) and Nr5a2 (right, blue) occupancy at regions bound by either of the two TFs. Esrrb/Nr5a2 peaks were ordered by decreasing Nr5a2 signal (mean Reads Per Million – RPM) and averaged every 20 regions. The full paired-end fragments were marked. **(G)** DNA sequence identified by de-novo motif discovery at all Esrrb/Nr5a2 target regions (top); note the 7th base can either be a T or a C. Boxplots showing Esrrb and Nr5a2 binding (RPM) at target regions containing tcaaggTca, tcaaggCca, both motifs or none, indicating the 7th base of the motif discriminates Esrrb (T) or Nr5a2 (C) preferential binding. **(H)** Heatmaps presenting Esrrb and Nr5a2 occupancy (RPM) at target regions ordered by descending Esrrb/Nr5a2 ratio (left), along the occurrence of the Esrrb or Nr5a2 motif as a function of the number of mismatches (mm; right).

To get initial indications of whether Esrrb and Nr5a2 could play overlapping functions in supporting pluripotency, we established their genome-wide binding profiles (Table S1). Cell lines in which a Flag peptide is knocked-in at the start of the endogenous Nr5a2 coding sequence were used (Fig S1), revealing extensive overlaps (Fig 1D,E). Possibly due to its lower expression, Nr5a2 binds fewer loci in ESCs, and almost invariably in association with Esrrb (Fig 1E,F). Yet, at common targets, preference for binding of one or the other factor was observed (arrowheads in Fig 1D, and Fig 1H). De novo motif discovery at regions bound by Esrrb or Nr5a2 found the canonical Esrrb consensus binding motif – TCAAGGTCA **(Festuccia et al., 2016)** – with the difference that either T or C could be accommodated at the 7th base of the motif (Fig 1G). To understand if the variation between T and C at this position could explain the preferential recruitment of Esrrb or Nr5a2 we analysed binding associated with each version of the consensus (TCAAGGTCA or TCAAGGCCA), revealing a preference of Esrrb for T and, more pronouncedly, of Nr5a2 for C. This was confirmed by ranking the regions bound by Esrrb and Nr5a2 by their preference for each factor, which revealed a strong bias of the Nr5a2 motif to regions with preferential Nr5a2 binding (Fig1H).

Altogether, we conclude that Esrrb and Nr5a2 display similar expression patterns in ESCs, where they bind at a common set of regulatory elements by virtue of highly similar, although exquisitely specific, DNA binding preferences.

### Esrrb and Nr5a2 are essential regulators of pluripotency

We have previously derived ESC lines in which endogenous Esrrb is knocked-out and Esrrb expression is rescued by a doxycycline (Dox) inducible transgene (EKOiE), such that upon Dox withdrawal the cells differentiate when cultured in FCS/LIF **(Festuccia et al., 2016)**. In these cells, we further disrupted the exon encoding the DNA binding domain of Nr5a2 at both endogenous alleles, to generate Nr5a2 null ESCs (EKOiE NrKO, Fig S1). EKOiE NrKO cells could be derived without special complications, indicating that in this context Nr5a2 is not by itself strictly required for the self-renewal of ESCs. Indeed, colony forming assays confirmed EKOiE NrKO cells self-renew despite showing increased spontaneous differentiation (Fig 2A,B). In comparison, the loss of Esrrb triggered by Dox withdrawal in EKOiE cells had more profound effects, resulting in the formation of few undifferentiated colonies, in line with previous results. Notably, the deletion of Nr5a2 exacerbated the effects of Esrrb loss of function, effectively ablating the formation of colonies containing undifferentiated cells 7 days after plating (Fig 2A,B). This suggests that in FCS/LIF Esrrb plays a preponderant role that is nevertheless further supported by Nr5a2, with clear additive effects. Substantiating these observations, the expression of pluripotency markers was more severely and consistently compromised two days after loss of both TFs, than after loss of Esrrb alone (Fig 2C). These trends were confirmed genome-wide: while Esrrb depletion leads to 707 differentially expressed genes, the concomitant loss of Nr5a2, which by itself deregulates 91 genes, is accompanied by extensive gene expression changes, with 1666 and 903 up- and down-regulated genes, respectively (FDR<0.01) (Fig 2D,S3A and Table S2). Moreover, we also observed that even though a small number of genes showed statistically significant changes in Nr5a2 KO, most of Esrrb targets displayed a concordant gene expression tendency, and vice versa (Fig 2D, S3B). Since genes activated and repressed by both factors are enriched in terms such as response to LIF (FDR=2.27e-24) and terms linked to differentiation (e.g. morphogenesis; FDR=2.32e-57), respectively, we conclude that Esrrb and Nr5a2 cooperate to support pluripotency in FCS/LIF.

**Fig. 2.**
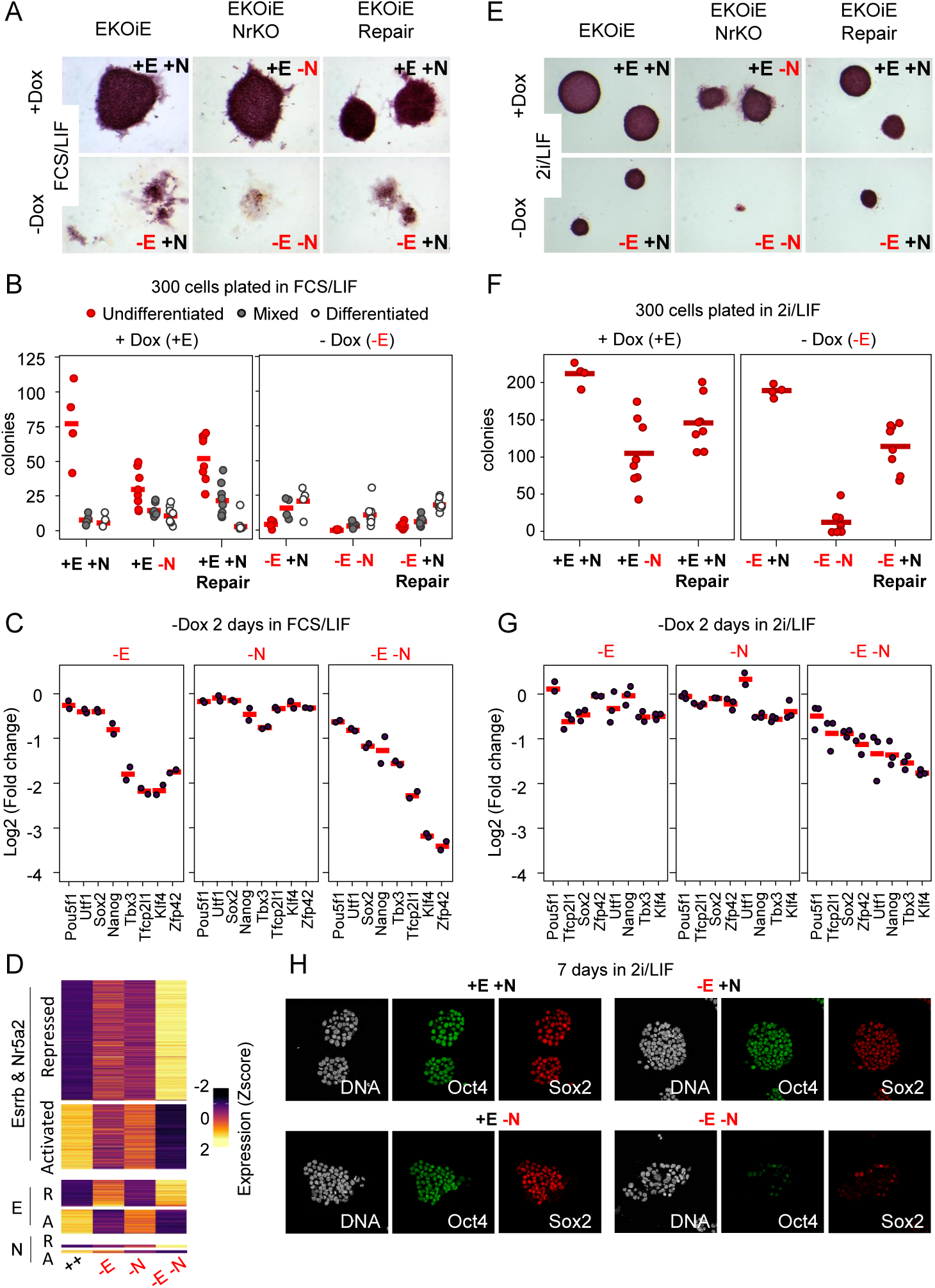
Esrrb and Nr5a2 are essential for ESCs self-renewal and pluripotency gene expression. **(A)** Representative images of alkaline phosphatase staining of colonies generated by the indicated ESC lines grown in FCS/LIF for 7 days. -E: loss of Esrrb expression; -N loss of Nr5a2 expression. **(B)** Quantification of the number of undifferentiated, mixed and differentiated colonies in the conditions described in (A). Each dot represents an independent experiment, the mean is marked by a red horizontal bar. **(C)** RNA-seq fold change of pluripotency gene expression in EKOiE ESCs two days after withdrawal of doxycycline (-E; left panel), in EKOiE NrKO ESCs compared to EKOiE cells (-N; middle panel) and in EKOiE NrKO ESCs two days after withdrawal of doxy-cycline, compared to EKOiE cells (-E-N; right panel). All cells were grown in FCS/LIF; each dot represents an independent experiment, the mean is marked by a red horizontal line. **(D)** Heatmaps showing Z-score of Transcripts Per Million (TPM) for genes identified as differentially expressed (absolute fold change >1.5; FDR < 0.01) in the conditions indicated on the left; -E, -N and -E-N as in (C). Cells were cultured in FCS/LIF. **(E-G)** Identical analyses to those described in (A-C) with cells cultured in 2i/LIF. **(H)** Immunofluorescence showing Oct4 and Sox2 expression in EKOiE and EKOiE NrKO cultured in 2i/LIF in the presence or absence of doxycycline for 7days. Note the virtually full depletion of Oct4 and Sox2 in the absence of both Esrrb and Nr5a2.

The extensive binding overlap between Esrrb and Nr5a2, their additive gene expression effects, and the severe phenotype of the deletion of both factors observed in ESCs cultured in FCS/LIF, prompted us to explore the effect of the concomitant loss of function in 2i/LIF, where the invalidation of Esrrb is compatible with the self-renewal of ESCs. Clonal plating and Dox withdrawal of EKOiE cells evidenced the detrimental effects of suppressing Esrrb expression, in line with previous reports **(Atlasi et al., 2019)**, yet confirming a non-essential function **(Martello et al., 2012)** (Fig 2E,F). Gene expression analysis further supports this conclusion, since acute Esrrb depletion in 2i/LIF results in reduced expression of both auxiliary and core pluripotency genes, in particular Klf4, Tbx3, Tfcp2l1 and Sox2 (Fig 2G). Yet expression of Oct4 is unaffected, and colonies of undifferentiated ESCs readily form after loss of Esrrb in 2i/LIF. Similarly, the loss of Nr5a2 had detrimental effects, reducing expression of pluripotency genes, the clonogenicity of ESCs and triggering spontaneous differentiation, while remaining overall tolerated **(Atlasi et al. 2019)**. In striking contrast, the concomitant depletion of Esrrb and Nr5a2 completely abolished the capacity of ESCs to self-renew, paralleling the observations made in FCS/LIF. The few colonies observed showed overt signs of morphological deterioration and included a large proportion of differentiated cells. These effects were specific, since repair of the disrupted Nr5a2 alleles rescued self-renewal (Fig 2E,F). In line, gene expression analysis revealed the collapse of pluripotency gene expression just two days after acute loss of Esrrb and Nr5a2 function (Fig 2G). Immunofluorescence 7 days after doxycycline withdrawal, also confirmed the near complete loss of Oct4, Sox2, Nanog and Klf4 expression in EKOiE NrKO ESCs (Fig 2H, S3C). These results establish Esrrb and Nr5a2 as a new set of essential regulators of pluripotency that act redundantly in ESCs to promote self-renewal.

### Loss of Esrrb and Nr5a2 triggers the collapse of the pluripotency network

The profound consequence of the loss of Esrrb and Nr5a2 function prompted us to investigate in more detail how these TF conjunctly control pluripotency TF binding, as represented by Oct4, Sox2 and Nanog, in ESCs cultured in 2i/LIF (Table S3). We identified 62,858 Esrrb binding regions in EKOiE cells grown in the presence of doxycycline (Fig S4A). As expected, binding was ablated 2 days after withdrawal of the drug. In contrast, the loss of Nr5a2 in EKOiE NrKO had no noticeable effect on Esrrb binding, suggesting a large independence in their binding activity (Fig S4B). In addition, we found that around 60% of the regions bound by Oct4, Sox2 and Nanog were also bound by Esrrb, in line with the notion that the pluripotency gene regulatory network is characterized by an extensively interconnected binding of TFs. Yet, around half of Esrrb binding events takes place at regions not targeted by any of the other TFs, as previously shown **(Festuccia et al., 2016)**. Importantly, however, maximal Esrrb, Oct4, Sox2 and Nanog binding was observed at regions of cobinding, suggesting a strong global cooperativity (Fig S4B). Strikingly, we observed that, at these common regulatory nodes, the loss of Esrrb and Nr5a2 leads to a global reduction of Oct4, Sox2 and Nanog occupancy (Fig 3A-C, S4C). While the reduced levels of Nanog and Sox2 protein 2 days after Nr5a2 and Esrrb depletion may partially contribute to these effects, Oct4 remained robustly expressed (Fig S4D). Moreover, the effect observed at Esrrb bound regions was significantly stronger than the reduction of binding observed at the ensemble of sites of pluripotency TF binding, irrespective of Esrrb occupancy (Fig S4C). Overall, more than 43%, 58% and 73% of co-bound regions displayed reduced Oct4, Sox2 and Nanog occupancy after depletion of Esrrb and Nr5a2, respectively. These effects are substantially more severe than those observed after depletion of Nanog in FCS/LIF **(Heurtier et al., 2019)**, and parallel in magnitude those observed after loss of Oct4, which was shown to reduce accessibility at 72% of its target enhancers **(King and Klose, 2017)**. We conclude that Esrrb and Nr5a2 are globally required to foster binding of Oct4, Sox2 and Nanog across thousands of regulatory regions.

**Fig. 3.**
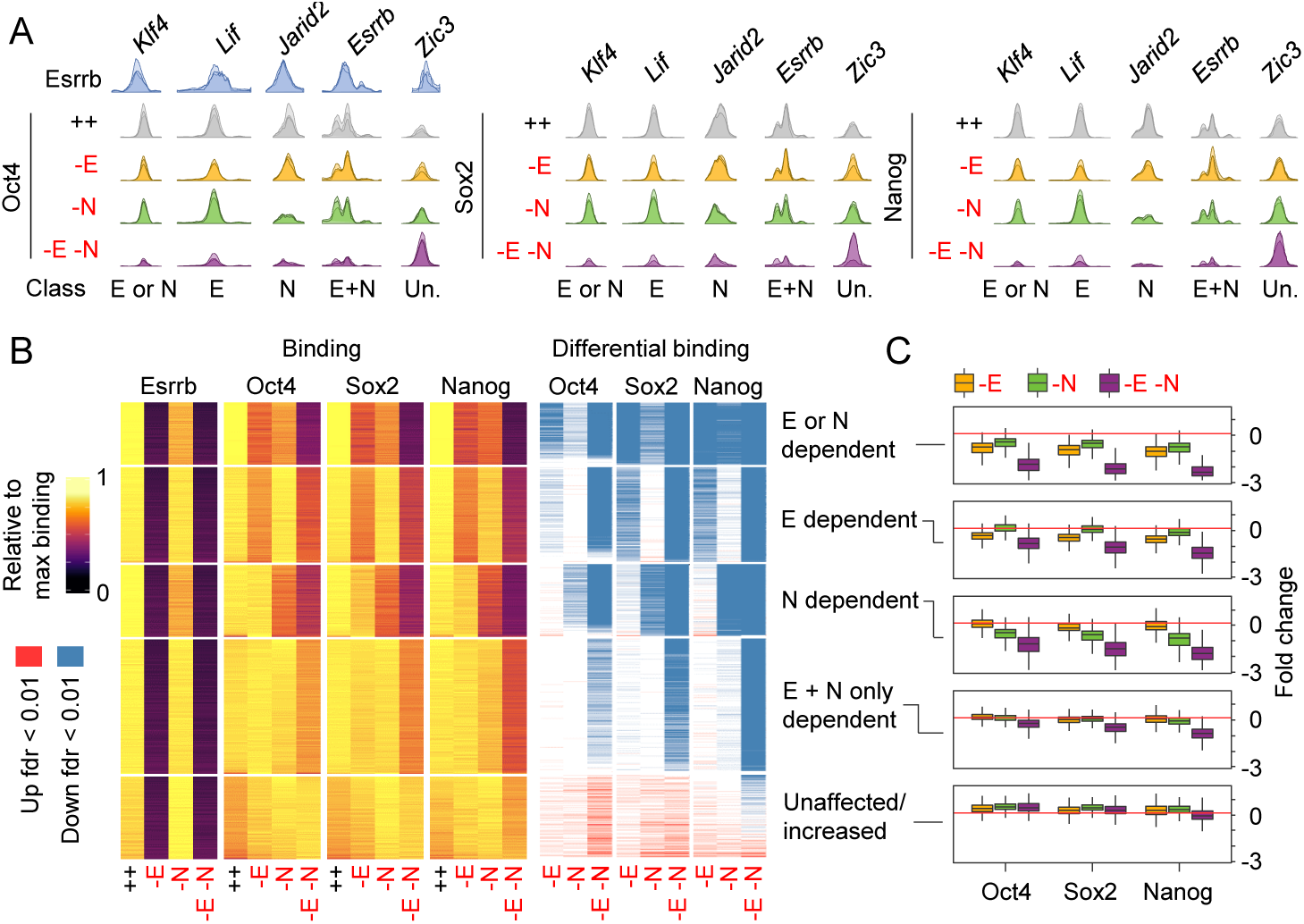
Esrrb and Nr5a2 control binding of Oct4, Sox2 and Nanog at thousands of regulatory elements. **(A)** ChIP-seq profiles of Esrrb, Oct4, Sox2 and Nanog binding at enhancers in proximity of the indicated genes in EKOiE and EKOiE NrKO cells cultured in 2i/LIF with or without doxycycline for two days, show-casing elements differentially affected by the depletion of Esrrb or Nr5a2. **(B)** Left, Binding: Heatmap showing normalized Esrrb, Oct4, Sox2 and Nanog binding levels (for each TF, the condition displaying maximal RPM is set to one) at 5 classes of regions identified by K-means clustering of Oct4, Sox2 and Nanog occupancy in EKOiE and EKOiE NrKO ESCs cultured 2 days in 2i/LIF in the presence or absence of doxycycline. Right, Differential binding calls for significantly increased (red) or decreased (blue) binding of each TF at the same set of regions (FDR < 0.05). The heatmap was simplified by averaging results of groups of 20 regions. **(C)** Fold change in Oct4, Sox2 and Nanog binding at the regulatory regions identified in B.

Next, we sought to further dissect the individual contributions of Esrrb and Nr5a2 to these effects, and identify affected and unaffected regulatory elements. Five major classes of regulatory regions were identified (Fig 3B). In the first group, we observed a significant loss of pluripotency TF binding upon the loss of either Esrrb or Nr5a2, and increased effects upon the dual loss. In the second and third groups all pluripotency factors lose binding in response to the loss of either Esrrb (group 2) or Nr5a2 (group 3), and show a more pronounced response after their combined depletion. In the fourth and largest group, the single depletions of Esrrb or Nr5a2 are relatively inconsequent and a significant reduction of pluripotency TF binding is observed with the double depletion. Finally, in the fifth group, we found regions that show unaltered or increased binding of Oct4 and Sox2 upon the loss of both Esrrb and Nr5a2. At these sites Esrrb and Nr5a2 oppose Oct4/Sox2 binding, which are also known to induce ESC differentiation when overexpressed **(Niwa et al., 2000)**. Clusters 1 to 4 tend to be located close to a sizable fraction of known pluripotency regulatory regions, such as the enhancers of Klf4, Nanog, Esrrb, Sox2 and Oct4. Indeed, they are collectively enriched in proximity of genes associated with pluripotency, such as response to LIF signalling (FDR<8.7e-7, all four clusters), blastocyst formation (FDR<2.5e-5, clusters 2 and 4) or stem cell population maintenance (FDR<5.1e-8, clusters 3 and 4). In contrast, cluster 5 is rather associated with genes linked to early epiblast differentiation, such as Grhl2/3, Zic2/3/5 and Fgf5, and neural fate (FDR=7.7e-7). These results indicate that Esrrb and Nr5a2, while displaying some specificity in function, tend to act cooperatively and are essential to maintain TF binding at the majority of regulatory elements they occupy, except at regions where the master pluripotency TFs Oct4/Sox2 are likely required to facilitate differentiation.

### Loss of pluripotency TF binding leads to major gene expression responses underlying the loss of self-renewal and the transition to early differentiation

We finally sought to determine which genes are deregulated in response the loss of Esrrb and Nr5a2 in ESC cultured in 2i/LIF. RNA-seq in EKOiE and EKOiE NrKO before or 2 days after doxycycline withdrawal identified genes differentially expressed after depletion of Nr5a2 (546 down-regulated, 515 upregulated) or Esrrb (343 down, 228 up), and highlighted that the targets of the two factors are largely overlapping (Fig 4A,B and Table S2). Nonetheless uniquely responsive genes could also be identified. In line with the pronounced effect on TF binding, loss of both Esrrb and Nr5a2 affected a higher number of genes than the individual depletion of either factor (1662 downregulated, 1779 upregulated). Of note, these transcripts show less pronounced but concordant changes after individual depletions, confirming that Esrrb and Nr5a2 largely cooperate in controlling gene expression (Fig 4A). We then measured the propensity of the 5 classes of regulatory regions identified above to be enriched in the vicinity of genes responding to the individual or combined depletion of Esrrb and Nr5a2. We observed a very strong statistical enrichment of genes activated by Esrrb and Nr5a2 around TF binding clusters 1 to 3, where the dual loss of Esrrb and Nr5a2 leads to a strong decrease of Oct4, Sox2 and Nanog binding (Fig 4C). Moreover, genes sensitive to Esrrb or Nr5a2 were also enriched in the vicinity of binding regions where either Esrrb or Nr5a2, respectively, are required for Oct4, Sox2 and Nanog binding, providing a direct functional support to our classification of regulatory regions. We also observed that regions showing a looser dependency on Esrrb or Nr5a2 displayed lower statistical enrichment around responsive genes. Finally, we noticed that genes repressed by Esrrb and/or Nr5a2 are associated with regions where Esrrb and/or Nr5a2 tend to oppose Oct4 and Sox2 binding. Altogether, this analysis suggests that Esrrb and Nr5a2 act more directly as activators than repressors, and that, to repress gene expression, both TFs often restrain Oct4/Sox2 binding.

**Fig. 4.**
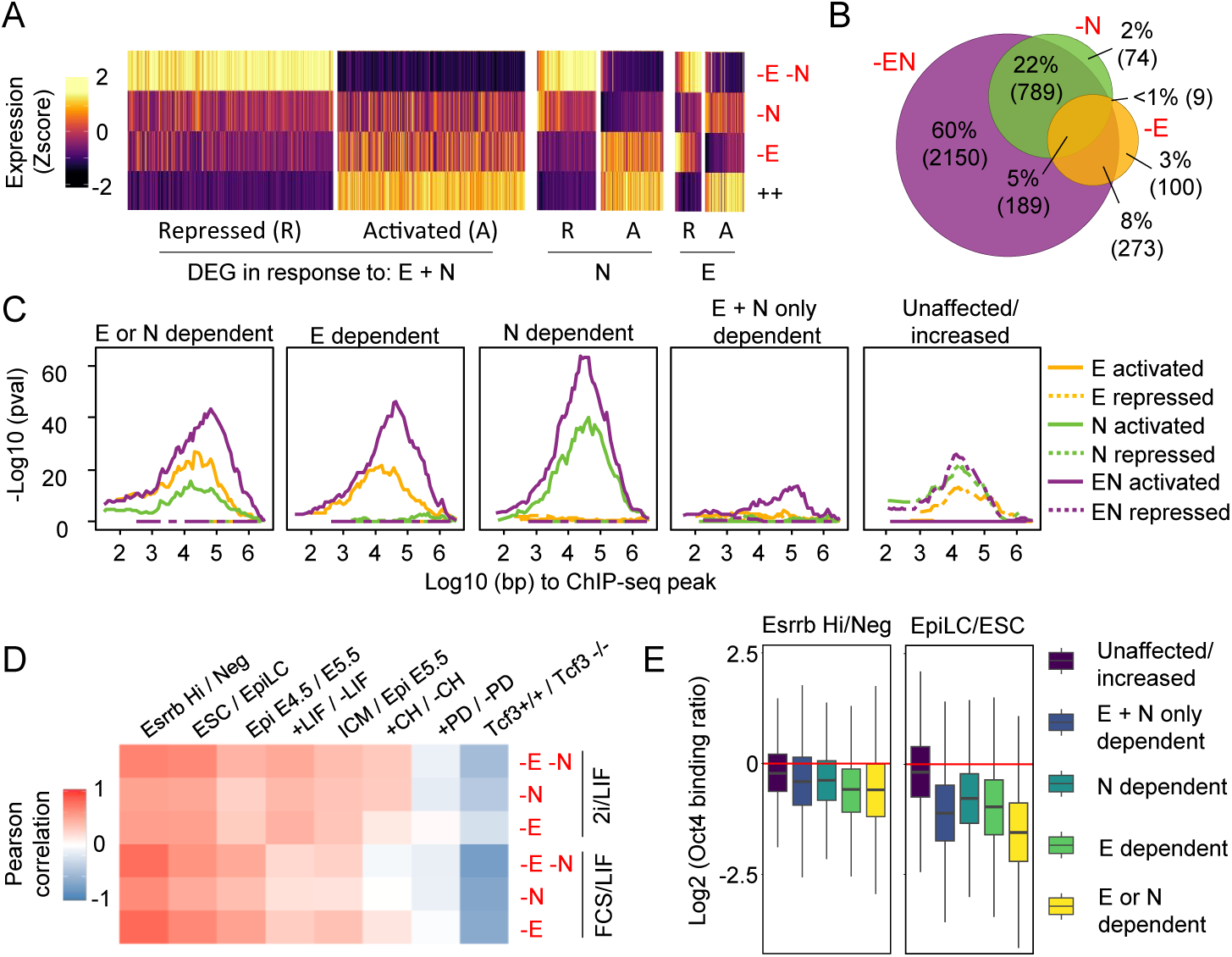
Esrrb and Nr5a2 maintain naïve pluripotency under the control of LIF and WNT signaling. **(A)** Heatmaps showing Z-score expression of genes significantly responding (FDR<0.01, absolute fold change>1.5) two days after inducing the loss of Esrrb, Nr5a2 or both TFs, in cells cultured in 2i/LIF, presented as in Figure 2D. **(B)** Venn diagram showing the overlap between genes responding to the depletion of Esrrb, Nr5a2 or both TFs. **(C)** Statistical enrichment (-log10 Hypergeometric right-tail p-value) of excess genes responsive to the depletion of Esrrb, Nr5a2, or both TFs as a function of the distance to the different classes of regions conjunctly bound by Esrrb, Oct4, Sox2 and Nanog identified in Figure 3B. **(D)** Heatmap displaying the Pearson correlation coefficient between the fold change in gene expression observed in this study after depletion of Esrrb, Nr5a2 or both TFs, and in the indicated conditions in published reports (see Methods for details). **(E)** Fold change in Oct4 binding between EsrrbHi and EsrrbNegative ESCs (as reanalyzed from Festuccia et al. EMBO J., 2018), or between ESC and Epiblast-like Stem Cells (EpiLC, as reanalyzed from Buecker et al. Cell Stem Cell, 2014), at the different classes of Esrrb and Nr5a2 responsive elements identified in Figure 3B.

We next explored the functional relevance of the gene expression changes mediated by Esrrb and Nr5a2. First, we observed that genes activated by Esrrb and Nr5a2 play a role in pluripotency regulation (response to LIF signalling, FDR=8.80e-11) and in energetic and metabolic processes (FDR=6.81e-6). Notably, when we compared the effect of the loss of Esrrb and Nr5a2 with the response to LIF stimulation in ESCs **(Martello et al., 2013)**, we observed a correlated response (Fig 4D, Pearson correlation coefficient 0.47). This prompted us to study more specifically the intersection of the activity of Esrrb/Nr5a2 with that of signalling pathways. In particular, both Esrrb and Nr5a2 have been suggested to act downstream of WNT **(Martello et al., 2012; Wagner et al., 2010)**. We thus set out to compare the effects of the loss of Tcf3 **(Yi et al., 2011)**, the main mediator of WNT activity in ESCs, and that of Esrrb and Nr5a2. Genes induced or downregulated in Tcf3-/- ESCs are respectively repressed or activated by Esrrb or Nr5a2 (Fig 4D, Pearson correlation coefficient Esrrb: −0.24, Nr5a2: −0.41). Notably, the concomitant loss of both Esrrb and Nr5a2 results in a higher correlation (−0.51). These results establish Esrrb and Nr5a2 as redundant mediators of the effects of WNT signalling in ESCs, possibly explaining why the individual deletion of either of these factors is tolerated in 2i/LIF. Altogether, these results suggest that Esrrb and Nr5a2 conjunctly act downstream of GSK3b inhibition, cooperating with LIF signalling to support ESC self-renewal.

We finally focused on genes upregulated after loss of both Esrrb and Nr5a2, and observed they are enriched for gene ontology terms linked to cell differentiation and developmental progression (FDR<7.34e-23). In agreement, we observed highly concordant transcriptional changes when we compared the depletion of Esrrb and Nr5a2 with those occurring during early differentiation in ESCs, both when occurring spontaneously in FCS/LIF **(Festuccia et al., 2018a)** and when directly driven from 2i/LIF cultures towards epiblast-like stem cells **(Buecker et al., 2014)** (Fig 4D, Pearson correlation coefficient 0.67, 0.64). This strongly indicates that Esrrb and Nr5a2 cooperate in restraining exit form naïve pluripotency. Moreover, since early differentiation is accompanied by a reorganization of Oct4 binding **(Buecker et al., 2014; Festuccia et al., 2018a)**, we compared the changes in Oct4 occupancy observed after loss of Esrrb and Nr5a2, with those previously reported. Regions showing increasing dependence on Esrrb and Nr5a2, also display a progressively more severe reduction in Oct4 binding either in spontaneously differentiating cells or during EpiLC conversion (Fig 4E). Reciprocally, enhancers maintaining or gaining Oct4 after Esrrb and Nr5a2 depletion (Group 5) are less affected during early differentiation. These correlations strongly suggest that Esrrb and Nr5a2 downregulation plays a causal role in triggering the transition from naïve to primed pluripotency.

Overall, our analysis establishes that Esrrb and Nr5a2 are required to assist TF binding at most of the regions they occupy: in their absence the pluripotency network collapses and the expression of pluripotency genes cannot be any longer supported by LIF and WNT. In addition, at a subset of developmental genes, Esrrb and Nr5a2 oppose Oct4 and Sox2 binding, possibly attenuating their differentiation-inducing effects. Through these two mechanisms, which are reminiscent of those observed for Nanog **(Heurtier et al., 2019)**, Esrrb and Nr5a2 mediate the fundamental function of maintaining the expression of developmental triggers in check. Since Esrrb and Nr5a2 are downregulated in the epiblast upon implantation, the function of both TFs in this context might be of developmental relevance. Supporting this, we observed that the genes displaying differential expression levels in early mouse embryos around the time of implantation, when the transition from naïve to prime pluripotency takes place **(Boroviak et al., 2015)**, are concordantly captured upon the combined loss of Esrrb and Nr5a2 in 2i/LIF (Fig 4D). These results therefore identify Esrrb and Nr5a2 as essential regulators, whose downregulation contributes to the ordered transition between pre and post-implantation pluripotency.

## Discussion

Our results identify a functional overlap between two members of the same family of orphan nuclear receptors, Esrrb and Nr5a2, in supporting pluripotency in ESCs. Similar redundant functions have been described for another family of auxiliary pluripotency TFs, KLFs **(Di Giammartino et al., 2019; Yamane et al., 2018)**. The individual deletion of Klf4, Klf2 and Klf5 is compatible with self-renewal in ESCs, but not the concomitant loss of all three TFs. While these results present analogies with our findings, these studies were not performed in conditions in which self-renewal is reinforced, as in 2i/LIF. It thus remains to be determined if KLF factors are strictly required for the maintenance of the undifferentiated state in ESCs. Intriguingly, it was shown that not all member of this family are able to support self-renewal, which is explained by variations in their zinc finger DNA binding domain **(Yamane et al., 2018)**. Of note, Nr5a2 and Esrrb are part of a group of orphan nuclear receptors that possess an extended DNA binding domain, which allows binding of these factors to DNA as monomers **(Gearhart et al., 2003; Solomon et al., 2005)**. In consequence, this class of nuclear receptors recognises a half-palindromic, extended, estrogen response element on DNA. The ability to access similar binding sites at regulatory regions emerges therefore as a common requirement for the redundancy between members of the same family of TFs. In this regard it is of note that, while we observed the ability of Esrrb and Nr5a2 to discriminate for the presence of T or C at the 7th base in their consensus binding sequence, the specificity in binding to one or the other version of the motif is nuanced, in particular for Esrrb. While the presence of an exact match to the Nr5a2 consensus is highly enriched in regions showing prevalent Nr5a2 binding, perfect Esrrb motifs preferentially appear in regions showing balanced occupancy by the two TFs. Considering the lower expression levels, this may indicate that high Nr5a2 binding requires either the presence of a strong Nr5a2 motif, or cooperativity driven by robust Esrrb occupancy. In addition to Esrrb and Nr5a2, at least two other classes of orphan receptors recognizing extended half-sites – Nr1d1 and Nr1d2 (Rev-Erb alpha and beta) and Nr4a1 (NgfIb) **(Harding and Lazar, 1993; Wilson et al., 1992)** – are expressed in ESCs at levels comparable to Nr5a2. It will be important to test the extent of the functional interactions of these TFs with Esrrb and Nr5a2 in supporting pluripotency in ESCs. KLF factors have been proposed as direct positive targets of STAT3, and therefore as important mediators of LIF signalling **(Niwa et al., 2009)**. Esrrb has instead been suggested to act in parallel to LIF, at least in 2i **(Martello et al., 2012)**. While these are likely not absolute distinctions, it is possible that Esrrb and Klf4 represent the two classes of factors through which LIF and WNT signalling prevalently input into the pluripotency network. This leaves open the question of which are the main mediators of ERK activity, which is the third signalling axis modulated by 2i/LIF. Amongst the pluripotency genes downregulated by ERK in ESCs, Nanog might perform a central function, given that ERK directly regulates expression **(Hamilton and Brickman, 2014)** and phosphorylates this TF **(Kim et al., 2014)**, and that Nanog contrasts the function of FGF/ERK signalling in inducing primitive endoderm specification during early development **(Chazaud et al., 2006; Frankenberg et al., 2011; Nichols et al., 2009)**. Nonetheless, Esrrb could also play an important part in mediating the effects of ERK inhibition: ERK has been shown to phosphorylate and control the activity of subunits of the Mediator complex **(Hamilton et al., 2019)**, that Esrrb deploys at enhancers of the pluripotency network to activate transcription **(Bell et al., 2020; Percharde et al., 2012)**. It remains to be addressed whether Nr5a2 displays a similar ability to interact with the transcriptional machinery in ESCs.

Our results question which TFs should be considered as “core” regulators of pluripotency. Oct4 and Sox2 are prominent amongst other pluripotency TFs, in that they perform essential functions both during development and in ESCs **(Avilion et al., 2003; Nichols et al., 1998)**. Another TF, Nanog, is required for epiblast specification **(Mitsui et al., 2003; Silva et al., 2009)**, but is not essential in ESCs, where it fine-tunes rather than enables self-renewal **(Chambers et al., 2003; Chambers et al., 2007)**. Here we show that Esrrb and Nr5a2 collectively play an essential role in ESCs. While detrimental, the loss of either factor alone does not fully compromise the maintenance of pluripotency **(Atlasi et al., 2019; Festuccia et al. 2012; Martello et al. 2012; Gu et al., 2005)**. In line, neither the loss of Esrrb, nor that of Nr5a2 results in developmental defects before implantation **(Gu et al., 2005; Labelle-Dumais et al., 2006; Luo et al., 1997)**. Esrrb deletion leads to defective placental development, and developmental arrest around E9.5, while Nr5a2 ablation results in gastrulation defects, and a severe phenotype emerges only at E7.5. In light of our results in ESCs, it will be now important to determine the effect of the concomitant loss of these TFs during early development. A clear requirement for the establishment of pluripotency would elevate Esrrb and Nr5a2 au pair with Oct4 and Sox2. Even then, while Esrrb and Nr5a2 are downregulated upon implantation, Oct4 and Sox2 continue playing an essential function in primed pluripotent cells **(Brons et al., 2007; Mulas et al., 2018; Osorno et al., 2012; Tesar et al., 2007)**. Our results could thus call for a distinction between core naïve activities, and global regulators of pluripotency. More directly, our results suggest a potential role for Esrrb and Nr5a2 in opposing the premature extinction of pluripotency in the naïve epiblast. Indeed, we show a clear analogy between the changes in gene expression and TF binding triggered by the loss of Esrrb and Nr5a2 and those observed during the conversion between naïve and primed pluripotency, both in culture and during development. This confirms previous results highlighting how, during the early stages of ESC differentiation, the spontaneous downregulation of Esrrb instates a transcriptional state that displays analogies to primed pluripotency **(Festuccia et al., 2018a)**. While the activity of other TFs – such as Foxd3, Otx2, and Grhl2 – have been proposed to play a determining role in rewiring TF binding during the dismantling of naïve pluripotency **(Buecker et al., 2014; Chen et al., 2018; Respuela et al., 2016; Yang et al., 2014)**, our results suggest that the loss of naïve TFs expression, and thus of cooperative interactions, plays an equally determinant role in driving the transition between pluripotent states.

Our results, extending previous reports, set the basis for the construction of a mechanistic framework that will allow to understand to what extent the activity of the pluripotency network depends on the function of its single components, and how, through these dependencies, it integrates the input of multiple signalling pathways to endow ESCs with essential developmental plasticity. The emerging picture suggests that the signalling axes controlling the pluripotency network conform to what have been proposed to constitute three general properties, or “habits”, of developmental pathways: activator insufficiency, cooperative action, and default repression **(Barolo and Posakony, 2002)**. No single TF responding to modulation of LIF, ERK or WNT signalling is in itself able to maintain ESC-specific gene expression, or coordinate its ordered rewiring during differentiation. Only the mutual dependency between TFs for accessing regulatory regions and activating gene expression ensures the robustness of the pluripotency network, while maintain its ability to be dismantled when single components are modulated in response to developmental cues, or constitutive repressive activities – such as that of Tcf3 on Esrrb – are no longer kept in check. In line, multiple mechanisms of cooperativity between pluripotency TFs have been documented: from the ability of Oct4 and Sox2 to bind DNA as heterodimers **(Remenyi et al., 2003; Tapia et al., 2015; Yuan et al., 1995)**, to the recognition by different members of the same class of TFs of related binding motifs – as for KLFs **(Yamane et al., 2018)** or Esrrb and Nr5a2 – to more indirect interaction mediated by the chromatin. In particular, Oct4 or Nanog have been reported to instate accessibility at target enhancers to foster TF binding **(Heurtier et al., 2019; King and Klose, 2017)**. It will be now important to determine how chromatin remodelers, in particular, SWI/SNF and NuRD complexes, which interacts with Esrrb in ESCs **(van den Berg et al., 2010)**, mediate the molecular activities we report here for Esrrb and Nr5a2. Moreover, it needs to be understood how the relative arrangement of TF binding sites at enhancers determines their dependency on Esrrb and Nr5a2 or on other pluripotency regulators.

Altogether, we report how Esrrb and Nr5a2, mediating the response to extrinsic cues, are essential in ESCs to instruct TF binding and maintain the wiring of the naïve pluripotency network. These results suggest that these two TFs are at the top of the hierarchy of gene regulation in ESCs, thereby opening new research avenues in the fields of developmental and stem cell biology, not only in mouse but also in humans. Indeed, the requirement of orphan nuclear receptors for the self-renewal of mouse ESCs stresses the need to understand their contribution to the maintenance of pluripotency in human ESC, especially in the less well characterized naïve state, where other nuclear receptors may be playing a preponderant role. By analogy to the functional overlap described between Esrra and Esrrg during somatic cell reprogramming and heart development **(Dufour et al., 2007; Kida et al., 2015)**, it is tempting to speculate that different combinations of nuclear orphan receptors may play a conserved role in pluripotency and during mammalian embryogenesis.

## Acknowledgements

This study was conceived, designed and initiated by N.F. during his stay at the MRC London Institute of Medical Sciences as a principal investigator. N.O performed computational analyses with help from A.C. A.D provided technical help. All experiments were performed by N.F, first at the LMS funded by an Imperial College Research Fellowship and an MRC Career Development Award, and then at the Institut Pasteur. N.F and P.N analysed the results and wrote the manuscript. P.N acknowledges the European Research Council (Erc-cog-2017, BIND), the Labex Revive (Investissement d’Avenir; ANR-10-LABX-73), the Institut Pasteur and the CNRS for funding.

## SUPPLEMENTARY INFORMATION

Four supplementary figures accompany this manuscript:

**Fig. S 1.**
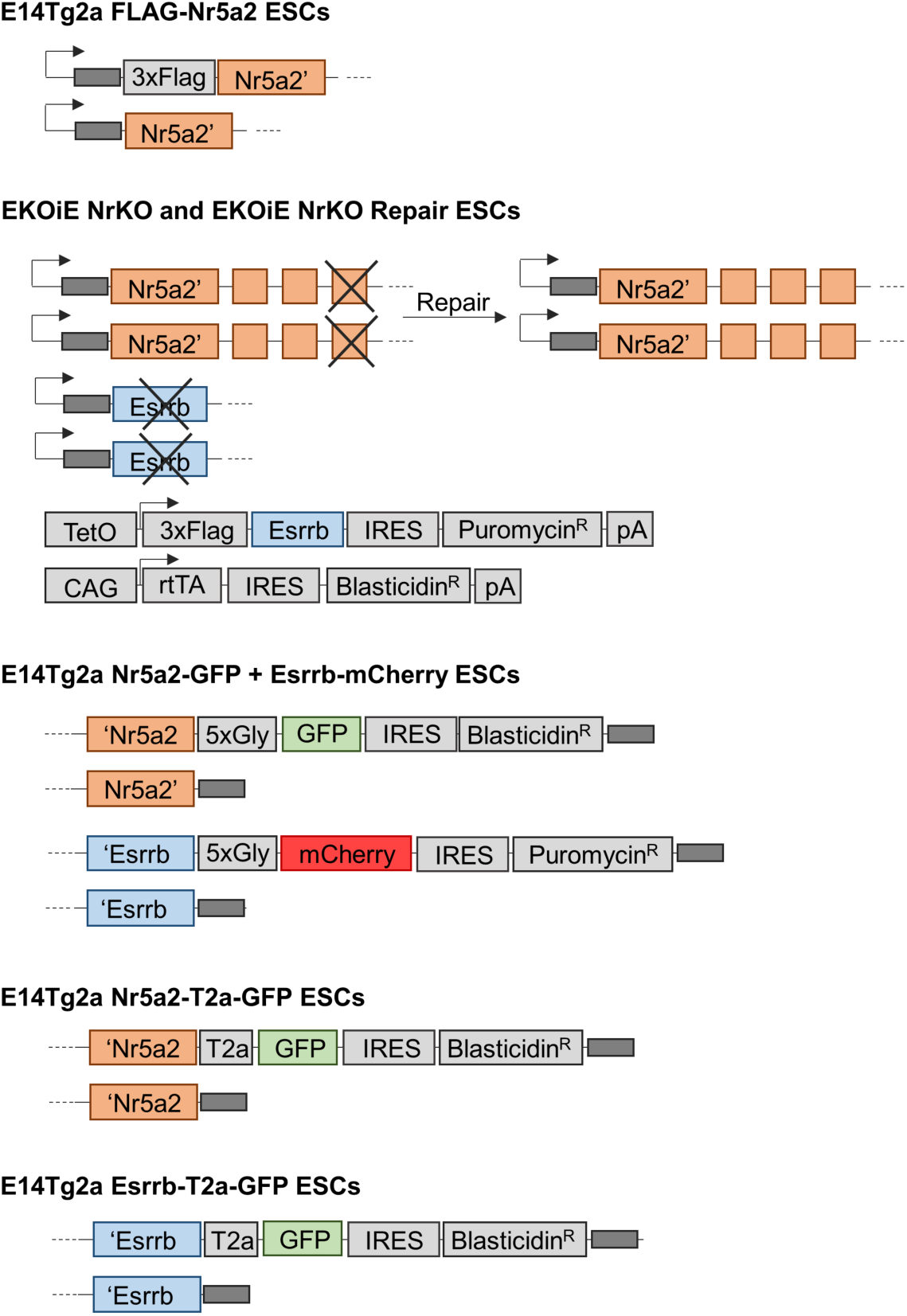
Cell lines used in this study.

**Fig. S 2.**
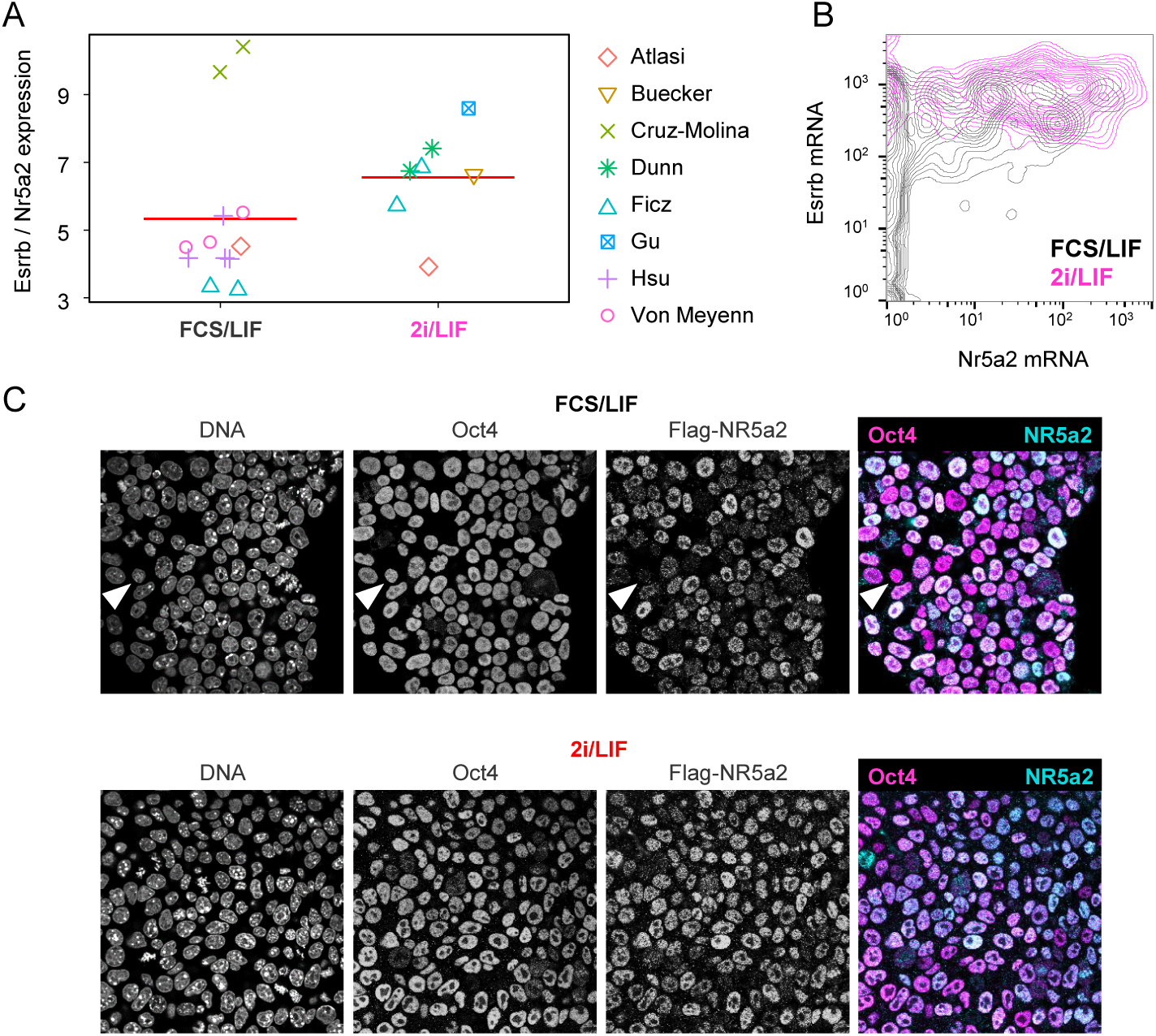
Additional information on Esrrb and Nr5a2 binding. **(A)** Plot indicating the ratio between Esrrb and NR5a2 gene expression levels in RNA-seq datasets from the indicated studies (See methods for a detailed list). Each dot represents an independent experiment, the red bars indicate the mean of values in each condition (FCS/LIF or 2i/LIF). **(B)** Density plot showing the distribution of Esrrb and Nr5a2 mRNA levels detected in single ESC cells cultured in FCS/LIF or 2i/LIF (data from Kolodziejczyk et al. Cell Stem Cell, 2015). **(C)** Confocal microscopy images showing Oct4 and Nr5a2 expression detected by immunofluorescence in Flag-Nr5a2 E14Tg2a ESC cultured in FCS/LIF or 2i/LIF. Note that Nr5a2 negative cells are prevalently observed in FCS/LIF (white arrowheads).

**Fig. S 3.**
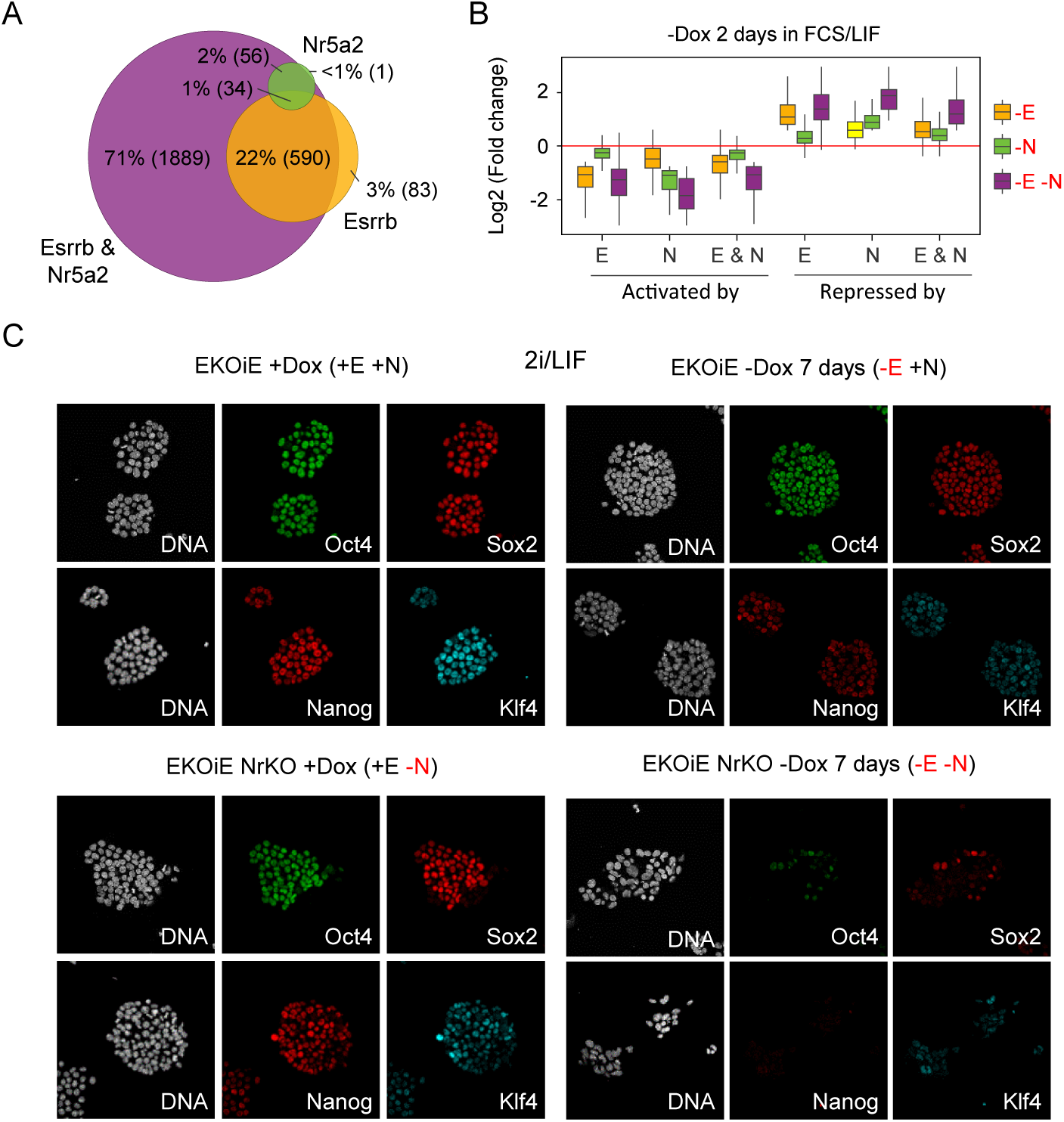
Additional information on the effects of the depletion of Esrrb and/or Nr5a2. **(A)** Venn diagram showing the overlap between genes responding to the depletion of Esrrb, Nr5a2 or both TFs. **(B)** Boxplots showing the fold change of gene expression of Esrrb, Nr5a2 and Esrrb/Nr5a2 responsive genes in EKOiE ESCs two days after withdrawal of doxycycline (-E; Orange), in EKOiE NrKO ESCs compared to EKOiE cells (-N; Green) and in EKOiE NrKO ESCs two days after withdrawal of doxycycline, compared to EKOiE cells (-E-N; Purple). All cells were grown in FCS/LIF. **(C)** Confocal microscopy images showing Oct4, Sox2, Nanog and Klf4 expression detected by immunofluorescence in EKOiE and EKOiE NrKO cultured in 2i/LIF in the presence or absence of doxycycline for 7 days.

**Fig. S 4.**
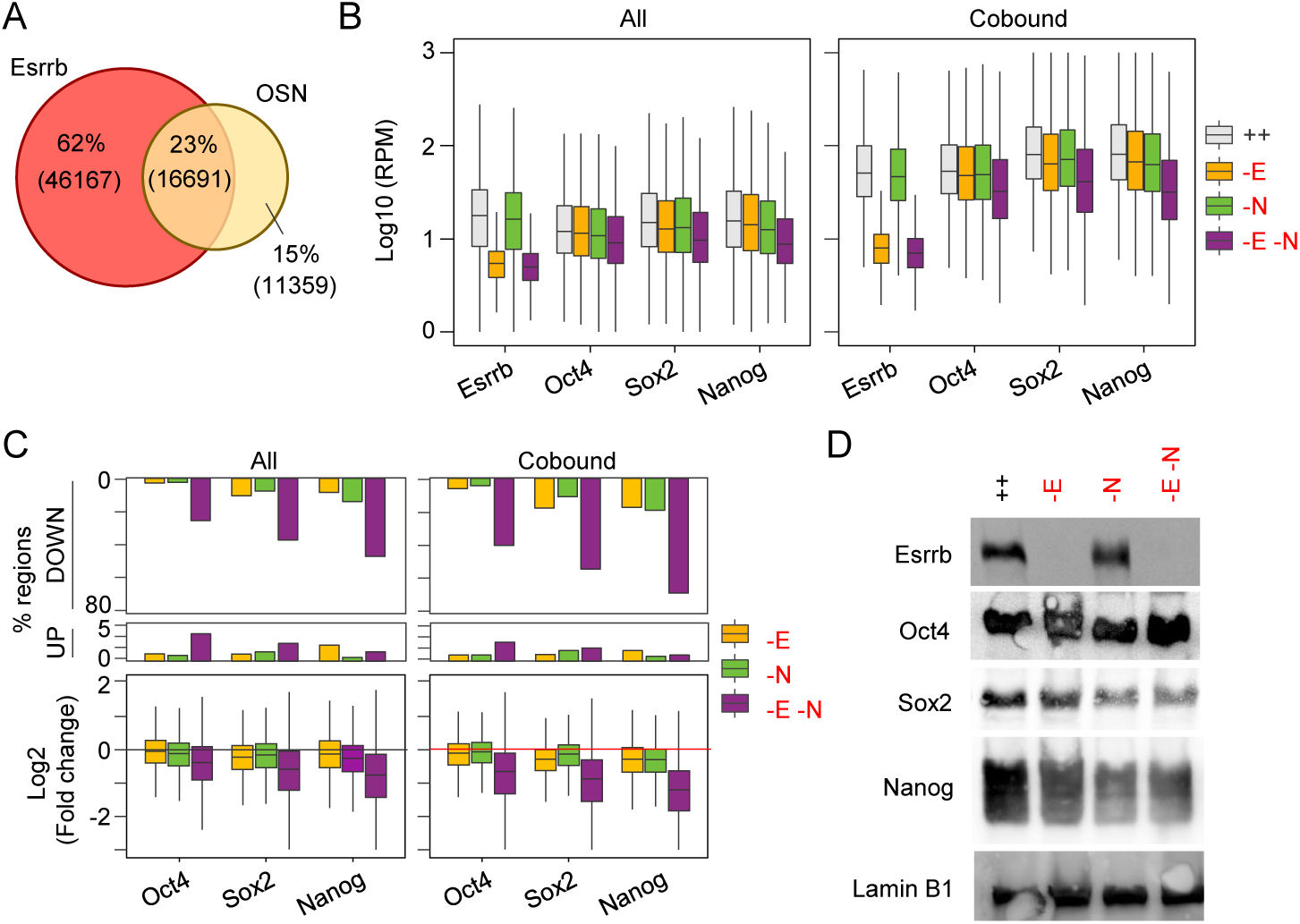
Additional information on pluripotency TF binding. **(A)** Venn diagram showing the intersection between regions called as bound by Esrrb or conjunctly by Oct4, Sox2 and Nanog (detected in all replicates, see Methods for details). **(B)** Boxplots showing Esrrb, Oct4, Sox2 and Nanog binding (RPM) at regions bound by each of the TFs, or at regions bound by all factors in conjunction. **(C)** Boxplot showing the fold change in Oct4, Sox2 and Nanog binding at regions bound by each of the TFs, or by the three factors in conjunction with Esrrb. **(D)** Western blot showing the levels of Esrrb, Oct4, Sox2, Nanog and Lamin B1 in EKOiE and EKOiE NrKO cultured in 2i/LIF in the presence or absence of doxycycline for 2 days.

Three Supplementary Tables are available online:

**Table S1:** Compendium of Esrrb/NR5a2 binding sites in FCS/LIF.

**Table S2:** All RNA-seq results are provided and annotated.

**Table S3:** Regions quantified for Esrrb, Oct4, Sox2 and Nanog binding in 2i/LIF

## References

Adachi, K., Kopp, W., Wu, G., Heising, S., Greber, B., Stehling, M., Arauzo-Bravo, M.J., Boerno, S.T., Timmermann, B., Vingron, M., et al. (2018). Esrrb Unlocks Silenced Enhancers for Reprogramming to Naive Pluripotency. Cell Stem Cell.

Atlasi, Y., Megchelenbrink, W., Peng, T., Habibi, E., Joshi, O., Wang, S.Y., Wang, C., Logie, C., Poser, I., Marks, H., et al. (2019). Epigenetic modulation of a hardwired 3D chromatin landscape in two naive states of pluripotency. Nat Cell Biol 21, 568–578.

Avilion, A.A., Nicolis, S.K., Pevny, L.H., Perez, L., Vivian, N., and Lovell-Badge, R. (2003). Multipotent cell lineages in early mouse development depend on SOX2 function. Genes Dev 17, 126–140.

Barolo, S., and Posakony, J.W. (2002). Three habits of highly effective signaling pathways: principles of transcriptional control by developmental cell signaling. Genes Dev 16, 1167–1181.

Bell, E., Curry, E.W., Megchelenbrink, W., Jouneau, L., Brochard, V., Tomaz, R.A., Mau, K.H.T., Atlasi, Y., de Souza, R.A., Marks, H., et al. (2020). Dynamic CpG methylation delineates subregions within super-enhancers selectively decommissioned at the exit from naive pluripotency. Nat Commun 11, 1112.

Boroviak, T., Loos, R., Lombard, P., Okahara, J., Behr, R., Sasaki, E., Nichols, J., Smith, A., and Bertone, P. (2015). Lineage-Specific Profiling Delineates the Emergence and Progression of Naive Pluripotency in Mammalian Embryogenesis. Dev Cell 35, 366–382.

Brons, I.G., Smithers, L.E., Trotter, M.W., Rugg-Gunn, P., Sun, B., Chuva de Sousa Lopes, S.M., Howlett, S.K., Clarkson, A., Ahrlund-Richter, L., Pedersen, R.A., et al. (2007). Derivation of pluripotent epiblast stem cells from mammalian embryos. Nature 448, 191–195.

Buecker, C., Srinivasan, R., Wu, Z., Calo, E., Acampora, D., Faial, T., Simeone, A., Tan, M., Swigut, T., and Wysocka, J. (2014). Reorganization of enhancer patterns in transition from naive to primed pluripotency. Cell Stem Cell 14, 838–853.

Chambers, I., Colby, D., Robertson, M., Nichols, J., Lee, S., Tweedie, S., and Smith, A. (2003). Functional expression cloning of Nanog, a pluripotency sustaining factor in embryonic stem cells. Cell 113, 643–655.

Chambers, I., Silva, J., Colby, D., Nichols, J., Nijmeijer, B., Robertson, M., Vrana, J., Jones, K., Grotewold, L., and Smith, A. (2007). Nanog safeguards pluripotency and mediates germline development. Nature 450, 1230–1234.

Chazaud, C., Yamanaka, Y., Pawson, T., and Rossant, J. (2006). Early lineage segregation between epiblast and primitive endoderm in mouse blastocysts through the Grb2-MAPK pathway. Dev Cell 10, 615–624.

Chen, A.F., Liu, A.J., Krishnakumar, R., Freimer, J.W., DeVeale, B., and Blelloch, R. (2018). GRHL2-Dependent Enhancer Switching Maintains a Pluripotent Stem Cell Transcriptional Subnetwork after Exit from Naive Pluripotency. Cell Stem Cell 23, 226–238 e224.

Di Giammartino, D.C., Kloetgen, A., Polyzos, A., Liu, Y., Kim, D., Murphy, D., Abuhashem, A., Cavaliere, P., Aronson, B., Shah, V., et al. (2019). KLF4 is involved in the organization and regulation of pluripotency-associated three-dimensional enhancer networks. Nat Cell Biol 21, 1179–1190.

Dufour, C.R., Wilson, B.J., Huss, J.M., Kelly, D.P., Alaynick, W.A., Downes, M., Evans, R.M., Blanchette, M., and Giguere, V. (2007). Genome-wide orchestration of cardiac functions by the orphan nuclear receptors ERRalpha and gamma. Cell Metab 5, 345–356.

Festuccia, N., Dubois, A., Vandormael-Pournin, S., Gallego Tejeda, E., Mouren, A., Bessonnard, S., Mueller, F., Proux, C., Cohen-Tannoudji, M., and Navarro, P. (2016). Mitotic binding of Esrrb marks key regulatory regions of the pluripotency network. Nat Cell Biol 18, 1139–1148.

Festuccia, N., Halbritter, F., Corsinotti, A., Gagliardi, A., Colby, D., Tomlinson, S.R., and Chambers, I. (2018a). Esrrb extinction triggers dismantling of naive pluripotency and marks commitment to differentiation. EMBO J 37.

Festuccia, N., Osorno, R., Halbritter, F., Karwacki-Neisius, V., Navarro, P., Colby, D., Wong, F., Yates, A., Tomlinson, S.R., and Chambers, I. (2012). Esrrb is a direct Nanog target gene that can substitute for Nanog function in pluripotent cells. Cell Stem Cell 11, 477–490.

Festuccia, N., Owens, N., and Navarro, P. (2018b). Esrrb, an estrogen-related receptor involved in early development, pluripotency, and reprogramming. FEBS Lett 592, 852–877.

Frankenberg, S., Gerbe, F., Bessonnard, S., Belville, C., Pouchin, P., Bardot, O., and Chazaud, C. (2011). Primitive endoderm differentiates via a three-step mechanism involving Nanog and RTK signaling. Dev Cell 21, 1005–1013.

Gearhart, M.D., Holmbeck, S.M., Evans, R.M., Dyson, H.J., and Wright, P.E. (2003). Monomeric complex of human orphan estrogen related receptor-2 with DNA: a pseudodimer interface mediates extended half-site recognition. J Mol Biol 327, 819–832.

Gu, P., Goodwin, B., Chung, A.C., Xu, X., Wheeler, D.A., Price, R.R., Galardi, C., Peng, L., Latour, A.M., Koller, B.H., et al. (2005). Orphan nuclear receptor LRH-1 is required to maintain Oct4 expression at the epiblast stage of embryonic development. Mol Cell Biol 25, 3492–3505.

Guo, G., and Smith, A. (2010). A genome-wide screen in EpiSCs identifies Nr5a nuclear receptors as potent inducers of ground state pluripotency. Development 137, 3185–3192.

Hamilton, W.B., and Brickman, J.M. (2014). Erk signaling suppresses embryonic stem cell self-renewal to specify endoderm. Cell Rep 9, 2056–2070.

Hamilton, W.B., Mosesson, Y., Monteiro, R.S., Emdal, K.B., Knudsen, T.E., Francavilla, C., Barkai, N., Olsen, J.V., and Brickman, J.M. (2019). Dynamic lineage priming is driven via direct enhancer regulation by ERK. Nature 575, 355–360.

Harding, H.P., and Lazar, M.A. (1993). The orphan receptor Rev-ErbA alpha activates transcription via a novel response element. Mol Cell Biol 13, 3113–3121.

Heng, J.C., Feng, B., Han, J., Jiang, J., Kraus, P., Ng, J.H., Orlov, Y.L., Huss, M., Yang, L., Lufkin, T., et al. (2010). The nuclear receptor Nr5a2 can replace Oct4 in the reprogramming of murine somatic cells to pluripotent cells. Cell Stem Cell 6, 167–174.

Heurtier, V., Owens, N., Gonzalez, I., Mueller, F., Proux, C., Mornico, D., Clerc, P., Dubois, A., and Navarro, P. (2019). The molecular logic of Nanog-induced self-renewal in mouse embryonic stem cells. Nat Commun 10, 1109.

Kida, Y.S., Kawamura, T., Wei, Z., Sogo, T., Jacinto, S., Shigeno, A., Kushige, H., Yoshihara, E., Liddle, C., Ecker, J.R., et al. (2015). ERRs Mediate a Metabolic Switch Required for Somatic Cell Reprogramming to Pluripotency. Cell Stem Cell 16, 547–555.

Kim, S.H., Kim, M.O., Cho, Y.Y., Yao, K., Kim, D.J., Jeong, C.H., Yu, D.H., Bae, K.B., Cho, E.J., Jung, S.K., et al. (2014). ERK1 phosphorylates Nanog to regulate protein stability and stem cell self-renewal. Stem Cell Res 13, 1–11.

King, H.W., and Klose, R.J. (2017). The pioneer factor OCT4 requires the chromatin remodeller BRG1 to support gene regulatory element function in mouse embryonic stem cells. Elife 6.

Labelle-Dumais, C., Jacob-Wagner, M., Pare, J.F., Belanger, L., and Dufort, D. (2006). Nuclear receptor NR5A2 is required for proper primitive streak morphogenesis. Dev Dyn 235, 3359–3369.

Luo, J., Sladek, R., Bader, J.A., Matthyssen, A., Rossant, J., and Giguere, V. (1997). Placental abnormalities in mouse embryos lacking the orphan nuclear receptor ERR-beta. Nature 388, 778–782.

Martello, G., Bertone, P., and Smith, A. (2013). Identification of the missing pluripotency mediator downstream of leukaemia inhibitory factor. EMBO J 32, 2561–2574.

Martello, G., Sugimoto, T., Diamanti, E., Joshi, A., Hannah, R., Ohtsuka, S., Gottgens, B., Niwa, H., and Smith, A. (2012). Esrrb is a pivotal target of the gsk3/tcf3 axis regulating embryonic stem cell self-renewal. Cell Stem Cell 11, 491–504.

Mitsui, K., Tokuzawa, Y., Itoh, H., Segawa, K., Murakami, M., Takahashi, K., Maruyama, M., Maeda, M., and Yamanaka, S. (2003). The homeoprotein Nanog is required for maintenance of pluripotency in mouse epiblast and ES cells. Cell 113, 631–642.

Mulas, C., Chia, G., Jones, K.A., Hodgson, A.C., Stirparo, G.G., and Nichols, J. (2018). Oct4 regulates the embryonic axis and coordinates exit from pluripotency and germ layer specification in the mouse embryo. Development 145.

Nichols, J., Silva, J., Roode, M., and Smith, A. (2009). Suppression of Erk signalling promotes ground state pluripotency in the mouse embryo. Development 136, 3215–3222.

Nichols, J., Zevnik, B., Anastassiadis, K., Niwa, H., Klewe-Nebenius, D., Chambers, I., Scholer, H., and Smith, A. (1998). Formation of pluripotent stem cells in the mammalian embryo depends on the POU transcription factor Oct4. Cell 95, 379–391.

Niwa, H., Miyazaki, J., and Smith, A.G. (2000). Quantitative expression of Oct-3/4 defines differentiation, dedifferentiation or self-renewal of ES cells. Nat Genet 24, 372–376.

Niwa, H., Ogawa, K., Shimosato, D., and Adachi, K. (2009). A parallel circuit of LIF signalling pathways maintains pluripotency of mouse ES cells. Nature 460, 118–122.

Osorno, R., Tsakiridis, A., Wong, F., Cambray, N., Economou, C., Wilkie, R., Blin, G., Scotting, P.J., Chambers, I., and Wilson, V. (2012). The developmental dismantling of pluripotency is reversed by ectopic Oct4 expression. Development 139, 2288–2298.

Percharde, M., Lavial, F., Ng, J.H., Kumar, V., Tomaz, R.A., Martin, N., Yeo, J.C., Gil, J., Prabhakar, S., Ng, H.H., et al. (2012). Ncoa3 functions as an essential Esrrb coactivator to sustain embryonic stem cell self-renewal and reprogramming. Genes Dev.

Remenyi, A., Lins, K., Nissen, L.J., Reinbold, R., Scholer, H.R., and Wilmanns, M. (2003). Crystal structure of a POU/HMG/DNA ternary complex suggests differential assembly of Oct4 and Sox2 on two enhancers. Genes Dev 17, 2048–2059.

Respuela, P., Nikolic, M., Tan, M., Frommolt, P., Zhao, Y., Wysocka, J., and Rada-Iglesias, A. (2016). Foxd3 Promotes Exit from Naive Pluripotency through Enhancer Decommissioning and Inhibits Germline Specification. Cell Stem Cell 18, 118–133.

Silva, J., Nichols, J., Theunissen, T.W., Guo, G., van Oosten, A.L., Barrandon, O., Wray, J., Yamanaka, S., Chambers, I., and Smith, A. (2009). Nanog is the gateway to the pluripotent ground state. Cell 138, 722–737.

Solomon, I.H., Hager, J.M., Safi, R., McDonnell, D.P., Redinbo, M.R., and Ortlund, E.A. (2005). Crystal structure of the human LRH-1 DBD-DNA complex reveals Ftz-F1 domain positioning is required for receptor activity. J Mol Biol 354, 1091–1102.

Tapia, N., MacCarthy, C., Esch, D., Gabriele Marthaler, A., Tiemann, U., Arauzo-Bravo, M.J., Jauch, R., Cojocaru, V., and Scholer, H.R. (2015). Dissecting the role of distinct OCT4-SOX2 heterodimer configurations in pluripotency. Sci Rep 5, 13533.

Tesar, P.J., Chenoweth, J.G., Brook, F.A., Davies, T.J., Evans, E.P., Mack, D.L., Gardner, R.L., and McKay, R.D. (2007). New cell lines from mouse epiblast share defining features with human embryonic stem cells. Nature 448, 196–199.

van den Berg, D.L., Snoek, T., Mullin, N.P., Yates, A., Bezstarosti, K., Demmers, J., Chambers, I., and Poot, R.A. (2010). An Oct4-centered protein interaction network in embryonic stem cells. Cell Stem Cell 6, 369–381.

Wagner, R.T., Xu, X., Yi, F., Merrill, B.J., and Cooney, A.J. (2010). Canonical Wnt/beta-catenin regulation of liver receptor homolog-1 mediates pluripotency gene expression. Stem Cells 28, 1794–1804.

Whyte, W.A., Orlando, D.A., Hnisz, D., Abraham, B.J., Lin, C.Y., Kagey, M.H., Rahl, P.B., Lee, T.I., and Young, R.A. (2013). Master transcription factors and mediator establish super-enhancers at key cell identity genes. Cell 153, 307–319.

Wilson, T.E., Paulsen, R.E., Padgett, K.A., and Milbrandt, J. (1992). Participation of non-zinc finger residues in DNA binding by two nuclear orphan receptors. Science 256, 107–110.

Yamane, M., Ohtsuka, S., Matsuura, K., Nakamura, A., and Niwa, H. (2018). Overlapping functions of Kruppel-like factor family members: targeting multiple transcription factors to maintain the naive pluripotency of mouse embryonic stem cells. Development 145.

Yang, S.H., Kalkan, T., Morissroe, C., Marks, H., Stunnenberg, H., Smith, A., and Sharrocks, A.D. (2014). Otx2 and Oct4 drive early enhancer activation during embryonic stem cell transition from naive pluripotency. Cell Rep 7, 1968–1981.

Yeo, J.C., and Ng, H.H. (2013). The transcriptional regulation of pluripotency. Cell Res 23, 20–32.

Yi, F., Pereira, L., Hoffman, J.A., Shy, B.R., Yuen, C.M., Liu, D.R., and Merrill, B.J. (2011). Opposing effects of Tcf3 and Tcf1 control Wnt stimulation of embryonic stem cell self-renewal. Nat Cell Biol 13, 762–770.

Ying, Q.L., Wray, J., Nichols, J., Batlle-Morera, L., Doble, B., Woodgett, J., Cohen, P., and Smith, A. (2008). The ground state of embryonic stem cell self-renewal. Nature 453, 519–523.

Yuan, H., Corbi, N., Basilico, C., and Dailey, L. (1995). Developmental-specific activity of the FGF-4 enhancer requires the synergistic action of Sox2 and Oct-3. Genes Dev 9, 2635–2645.

