## Supplementary material for "The combined action of Esrrb and Nr5a2 is essential for naïve pluripotency": Methods

**Contents:**

**1/ Culture and Generation of ES cells.** **page 2**
*1a/ General culture conditions.*
*1b/ Derivation of Esrrb-T2a-GFP, Nr5a2-T2a-GFP and Nr5a2-GFP+Esrrb-mCherry ESCs.*
*1c/ Derivation of FLAG-Nr5a2 ESCs.*
*1d/ Derivation of EKOiE NrKO and EKOiE NrKO Repair ESCs.*
*1e/ Alkaline phosphatase staining.*

**2/ Flow Cytometry.** **page 3**

**3/ Imaging.** **page 3**
*3a/ Immunofluorescence on fixed cells.*
*3b/ Live-cell imaging.*
*3c/ Imaging of alkaline phosphatase stainings.*

**4/ TF binding analysis by ChIP-seq.** **page 4**
*4a/ Chromatin immunoprecipitation (ChIP).*
*4b/ Library preparation.*

**5/ RNA-seq.** **page 5**

**6/ Western Blot.** **page 6**

**7/ Computational Methods.** **page 6**
*7a/ Data and availability.*
*7b/ ChIP-seq analyses.*
*7c/ RNA-seq analyses.*
*7d/ Comparisons to published datasets.*

**Supplementary references** **page 9**

### 1/ Culture and Generation of ES cells.

#### *1a/ General culture conditions.*

ES cells were cultured on 0.1% gelatine (SIGMA, Cat# G1890-100G) in DMEM + GlutaMax-I (Gibco, Cat# 31966-021), 10% FCS (Gibco, Cat# 10270-098), 100 µM 2-mercaptoethanol (Gibco, Cat# 31350-010), 1× MEM non-essential amino acids (Gibco, Cat# 1140-035) and 10 ng×ml$^{-1}$ recombinant LIF (MILTENYI BIOTEC, Cat# 130-099-895) (Referred to as FCS/LIF). Cells were passaged 1:10 every 2–3 days. When indicated, cells were grown in 2i-containing medium (1µM PD0325901 and 3 µM CHIR99021; Axon Medchem Bv) - 0.5× DMEM/F12 (Gibco, Cat# 31331093), 0.5×Neurobasal (Gibco, Cat# 21103049), 0.5% N2 supplement 100× (Gibco, Cat# 17502048), 1% B27 supplement 50× (Gibco, Cat# 17504044), 10µg/mL Insulin (Sigma, Cat# I1882-100MG), 2 mM L-Glutamine (Invitrogen, Cat# 91139), 0.05% BSA (Sigma, Cat# A3311-10G), 100 µM 2-mercaptoethanol (Gibco, Cat# 31350-010), 10 ng/ml recombinant LIF (MILTENYI BIOTEC, Cat# 130-099-895) (Referred to as 2i/LIF).

#### *1b/ Derivation of Esrrb-T2a-GFP, Nr5a2-T2a-GFP and Nr5a2-GFP + Esrrb-mCherry ESCs.*

E14Tg2a were nucleofected - using an Amaxa Nucleofector II with a Mouse ES Cell Kit (LONZA, Cat# VPH-1001) and selecting programme A23 - with 3µg of a linearised targeting vector designed to insert a T2a or 5×Gly linker – Fluorescent protein – IRES Blasticidin$^{R}$ cassette at the stop codon of *Esrrb* (ENSEMBL transcript Esrrb-206) or *Nr5a2* (ENSEMBL transcript Nr5a2-205) and 1µg of pU6_CBh-Cas9-T2A-mCherry (Addgene no. 64324) driving expression of the gRNAs 5′- TGTGCTGGGCCATCACACCT -3′ for Esrrb or 5′- GCTTCCAGGGGTGGGGACTT -3′ for Nr5a2. 48 hours later Blasticidin selection was added (5µg×ml$^{-1}$ ) and after 2 weeks single colonies were picked, expanded, and correctly targeted cells identified by PCR on genomic DNA followed by sequencing. An identical procedure was followed to derive Nr5a2-GFP+Esrrb-mCherry ESCs from heterozygous Nr5a2-GFP cells that were further nucleofected with 3µg of a linearised targeting vector designed to insert a 5×Gly linker – mCherry – IRES Puromycin$^{R}$ cassette at the stop codon of *Esrrb and* 1µg of pU6_CBh-Cas9-T2A-mCherry driving expression of the gRNA 5′- TGTGCTGGGCCATCACACCT -3′.

#### *1c/ Derivation of FLAG-Nr5a2 ESCs.*

E14Tg2a were nucleofected (See 1b) with 3µg of a linearised targeting vector designed to insert a LoxP – Puromycin$^{R}$ – LoxP – 3xFLAG – Gly5 – cassette at the start codon of *Nr5a2* (ENSEMBL transcript Nr5a2-205) and 1µg of a pU6_CBh-Cas9-T2A-mCherry plasmid driving expression of the gRNA 5′-CCACTTTGGGCAGCATGACA -3′. 48 hours later selection was added and after 1 week resistant cells were further nucleofected with 3µg of a plasmid driving expression of the Cre recombinase, and plated at clonal density. 2 weeks later , single colonies were picked, expanded, and correctly targeted cells, which had excised the Puromycin$^{R}$ cassette, identified by PCR on genomic DNA and by immunofluorescence with anti-Flag mouse monoclonal (M2 clone - Sigma Cat# F3165). Three heterozygous targeted clones (clones 2, 17 and 22) were selected for further experiments.

#### *1d/ Derivation of EKOiE NrKO and EKOiE NrKO Repair ESCs.*

EKOiE ESCs (Festuccia et al. 2016) were nucleofected (See 1b) with 3µg of an equimolar pool of 2 pU6_CBh-Cas9-T2A-mCherry plasmids driving expression of the gRNAs 5′- CCTCACGTGCGAAAGCTGCA -3′ or 5′- GACACTTTATCGCCACACAC -3′ to induce double strand breaks into the 4$^{th}$ Exon of *Nr5a2* (Nr5a2-205). 48 hours later single mCherry positive cells were FACS sorted into individual wells of a 96-well plate and expanded. Correctly targeted cells were identified by PCR on genomic DNA and sequencing. Two clones bearing homozygous deletions in exon 4 (clone 2; 53 bp deletion; mm10_Chr1:136944963-136945015) or (clone 4; 52 bp deletion; mm10_Chr1:136944963-136945014) of the Nr5a2 gene transcript (ENSEMBL Nr5a2-205) were selected for further experiments. Clone 4 EKOiE NrKO ESCs were further nucleofected with 1µg of a pU6_CBh-Cas9-T2A-mCherry plasmid driving expression of the gRNA 5′- GAAAAGGCAAACCTTGCCAC -3′ specific to the disrupted Nr5a2 alleles, and 7 µg of a 300bp-long repair template obtained by PCR from E14Tg2a genomic DNA (primers 5′- TCAGCAATGCTTTTCAGTGCAG -3′ and 5′- TGACACTTCCCCCACCTCAC -3′). 48 hours later single mCherry positive cells were FACS sorted into individual wells of a 96-well plate and expanded. Correctly repaired cells were identified by PCR on genomic DNA and sequencing. Two homozygously repaired clones (clone 4.2 and 16.1) were selected for further experiments.

***1e/ Alkaline phosphatase staining.***
300 EKOiE, EKOiE Nr5a2 KO (c2 and c4), and EKOiE Nr5a2 KO Repair ESCs (c 4.2 and 16.1) were plated in single wells of 6-well plates coated overnight with poly-L-ornithine 0.01% (Sigma, Cat# P4957) at 4 °C, washed and coated 2 h with laminin (Millipore, Cat# CC095) 10 $\mu g \times ml^{-1}$ in PBS. After culture in FCS/LIF or 2i/LIF media in presence or absence of 1μg/ml doxycycline for 7 days cells were fixed and stained using an alkaline phosphatase staining kit (Sigma Aldrich, Cat # 86R-1KT) according to the manufacturer's instructions.

**2/ Flow cytometry.**

E14Tg2a, Esrrb-T2a-GFP or Nr5a2-T2a-GFP ESCs were plated at low density (2000 cells/$cm^2$) in FCS/LIF medium and cultured for 3 days before analysis. After trypsinisation, cells were resuspended in FCS/LIF without phenol-red, and analysed using a LSR II flow cytometer system or a Luminex Image Stream MK2 instrument with a 60x magnification objective. Data was analysed using the FlowJo software suite.

**3/ Imaging.**

***3a/ Immunofluorescence on fixed cells.***
Cells were plated on IBIDI hitreat plates coated overnight with poly-L-ornithine 0.01% (Sigma, Cat# P4957) at 4 °C, washed and coated 2h with laminin (Millipore, Cat# CC095) 10 μg/ml in PBS. Fixation was performed for 10′ in 1% formaldehyde (Thermo, Cat# 28908) at room temperature. After washing the cells twice in PBS, they were permeabilised with PBS/0.1% v/v Triton X-100 supplemented with 3% of donkey serum (Sigma, Cat# D9663) for 30 min at room temperature. Primary antibodies (diluted in PBS with donkey serum 3%) were applied 2h at room temperature or overnight at 4°C in a volume of 1ml per dish. After three washes in PBS/0.1% Triton X-100, secondary antibodies (2μg/ml in PBS with donkey serum 3%) were applied for 2h at room temperature. Cells were washed three times in PBS/0.1% v/v Triton X-100, nuclei counterstained with 4',6-diamidino-2-phenylindole (DAPI; Sigma, Cat# D9542), and imaged with an inverted Leica SP8 confocal microscope using a 40X oil immersion objective. Acquisition was performed using the LASX acquisition software suite. Primary antibodies were used at the following concentrations:

- 0.3μg/ml anti-Nanog rabbit polyclonal (Cosmobio, Cat# REC-RCAB001P)
- 0.4μg/ml anti-Oct4 mouse monoclonal - clone C10 (Santa Cruz Biotechnology, Cat # sc-5279), for staining in combination with anti-Sox2
- 1μg/ml anti-Oct4 rabbit polyclonal (Abcam, Cat # ab19857), for staining in combination with anti-Flag
- 1 :500 anti-Sox2 rabbit polyclonal (Active Motif, Cat # 39843)
- 1μg/ml anti-Klf4 goat polyclonal (R&D, Cat # AF3158)
- 1 μg/ml anti-Flag mouse monoclonal (Sigma, Cat# F3165)

Secondary antibodies: Alexa Fluor 594 AffiniPure Donkey Anti-Rabbit IgG (H+L) (Jackson ImmunoResearch, Cat #711-585-152); Alexa Fluor 488 AffiniPure Donkey Anti-Mouse IgG (H+L) (Jackson ImmunoResearch, Cat #715-545-150); Alexa Fluor 647 AffiniPure Donkey Anti-Goat IgG (H+L) (Jackson ImmunoResearch, Cat # 705-605-147).

***3b/ Live-cell imaging.***

Nr5a2-GFP+Esrrb-mCherry ESCs grown in FCS/LIF or 2i/LIF in IBIDI plates as described above were incubated with 500 nM Hoechst-33342 for 20 min before imaging. During imaging the cells were kept at 37 °C in a humidified atmosphere (7% $CO_2$). Image on single focal planes were acquired with a 63X oil immersion objective on an inverted LSM800 confocal Zeiss microscope, using the ZEN Blue acquisition software suite.

***3c/ Imaging of alkaline phosphatase stainings.***

Representative images of alkaline phosphatase stained colonies formed by EKOiE, EKOiE Nr5a2 KO, and EKOiE Nr5a2 KO repair ESCs were acquired using a Zeiss Discovery V8 Stereo microscope, and the ZEN blue software suite.

**4/ TF binding analysis by ChIP-seq.**

***4a/ Chromatin immunoprecipitation (ChIP).***

After trypsinisation, $10^7$ ES cells were crosslinked in 2 ml of freshly prepared PBS-DSG 2 mM at pH 7.0 (Sigma, Cat# 80424-5 mg) for 50 min at room temperature with occasional shaking. After pelleting and washing once in PBS, cells were incubated for 10 min in 2 ml PBS 1% formaldehyde (Thermo, Cat# 28908). Crosslinking was stopped with 0.125 mM glycine for 5 min at room temperature. Cells were pelleted and washed with ice-cold PBS. Cells were resuspended in 2 ml of swelling buffer (25 mM Hepes pH 7.95, 10 mM KCl, 10 mM EDTA) freshly supplemented with 1× protease inhibitor cocktail (PIC-Roche, Cat# 04 693 116 001) and 0.5% IGEPAL. After 30 min on ice, the suspension was passed 40 times in a dounce homogenizer. Cells were then centrifuged and resuspended in 300 µl of TSE150 (0.1% SDS, 1% Triton X-100, 2 mM EDTA, 20 mM Tris-HCl pH8, 150 mM NaCl) buffer, freshly supplemented with 1× PIC. Samples were sonicated in 1.5 ml tubes (Diagenode) using a Bioruptor Pico (Diagenode) for 7 cycles divided into 30 s ON–30 s OFF sub-cycles at maximum power, in circulating ice-cold water. After centrifugation (30 min, full speed, 4 °C), the supernatant was either used immediately for immunoprecipitation or stored at −80 °C until use, generally within the month. Five microliters were used to quantify the chromatin concentration and check DNA size (typically 200–350 bp). Chromatin from $10^7$ cells was used for each ChIP-seq after pre-clearing it for 3 hours rotating on-wheel at 4 °C in 300 µl of TSE150 containing 50 µl of protein G Sepharose beads (Sigma, Cat# P3296-5 ML) 50% slurry, previously blocked with BSA (500 µg×ml$^{-1}$; Roche, Cat# 5931665103) and yeast tRNA (1 µg×ml$^{-1}$; Invitrogen, Cat# AM7119). Immunoprecipitations with anti-Esrrb mouse monoclonal (1µg per 2x$10^6$cells, Perseus Proteomics, Cat# H6-705-00), anti-Nanog rabbit polyclonal (0.6 µg per 2x$10^6$cells, Cosmobio, Cat# REC-RCAB001P); anti-Oct4 rabbit polyclonal (1 µg per 2x$10^6$cells, Abcam Cat # ab19857); anti-Sox2 rabbit polyclonal (1 µl per 2x$10^6$cells – concentration not specified, Active Motif Cat# 39844); anti-Flag mouse monoclonal (1 µg per 2x$10^6$cells Sigma Cat# F3165) antibodies, were performed overnight rotating on-wheel at 4 °C in 500 µl of TSE150. 20 µl were set apart for input DNA extraction and precipitation. 25 µl of blocked protein G beads 50% slurry was added for 4 h rotating on-wheel at 4 °C. Beads were pelleted and washed for 5 min rotating on-wheel at room temperature with 1 ml of buffer in the following order: 3 × TSE150, 1 × TSE500 (as TSE150 but 500 mM NaCl), 1× washing buffer (10 mM Tris-HCl pH8, 0.25M LiCl, 0.5% NP-40, 0.5% Na-deoxycholate, 1 mM EDTA), and 2 × TE (10 mM Tris-HCl pH8, 1 mM EDTA). Elution was performed in 100 µl of elution buffer (1% SDS, 10 mM EDTA, 50 mM Tris-HCl pH 8) for 15 min at 65 °C after vigorous vortexing. Eluates were collected after centrifugation and beads rinsed in 150 µl of TE-SDS1%. After centrifugation, the supernatant was pooled with the corresponding first eluate. For both immunoprecipitated and input chromatin, the crosslinking was reversed overnight at 65 °C, followed by proteinase K treatment, phenol/chloroform extraction and ethanol precipitation.

**4b/ *Library preparation.***

*SPRI Bead preparation:* 1 ml Sera-Mag™ Magnetic SpeedBeads™, carboxylated, 1 µm, 3 EDAC/PA5 (GE Healthcare Life Sciences, Cat# 65152105050250) were washed 3 times with a TE-Tween solution (10 mM Tris HCl pH 8, 1 mM EDTA, 0.05% Tween 20, pH 8.0) and resuspended in TE-Tween-20% PEG 8000 solution (10 mM Tris HCl pH 8, 1 mM EDTA, 0.05% Tween 20, pH 8.0).

*End repair:* Precipitated DNA was resuspended in 37.5µl of water and mixed with 2µl of 10mM dNTPs, 5µl of NEB T4 ligase buffer, 2.5µl of NEB T4 polymerase (Cat# M0203L), 0.5µl of NEB Klenow polymerase (Cat# M0210L) and 2.5µl of NEB T4 PNK (Cat# M0201L). Samples were incubated 30min at 20°C in a thermocycler. DNA was purified with SPRI beads: 90µl of SPRI bead suspension and 50µl isopropanol were added and samples transferred to a 96 well plate. After incubating for 5 min, the plate was put on a 96S Super Ring Magnet (Alpaqua, Cat# A001322), beads were allowed to separate completely, and the supernatant removed without disrupting the bead pellet. Beads were washed twice with 200µl of 70% Ethanol and the supernatant completely removed. DNA was eluted in 21µl of water.

*A-Tailing:* 20µl of sample were mixed with 2.5µl of NEB Buffer #2, 1µl of 5mM dATP, 1.5µl of NEB Klenow 3'-5' exo minus (Cat# M0212L), and incubated at 37°C for 30min in a thermocycler. DNA was purified with SPRI beads as before, but using a volume of 45µl of beads and 25µl isopropanol. DNA was elute in 20µl of water.

*Adaptor ligation:* 19µl of sample were mixed with 2.5µl of NEB T4 ligase buffer, 1.25µl of a 0.2µM solution of annealed adapters, and 2.5µl of NEB concentrated T4 ligase (Cat# M0202M) and incubated overnight at 16°C. DNA was purified with SPRI beads as before, but using a volume of 35µl of beads and no isopropanol, eluting in 20 µl of water. Adaptors were designed in house based on the structure of Illumina TruSeq™ indexed and forked adapters, modified to include extended indexes. 22.5µl each of 40µM ssDNA Barcoded and Universal adaptor solutions were mixed with 5µl of NEB buffer 2, and annealed in a thermocycler.

Barcoded adaptor:

5'P-GATCGGAAGAGCACACGTCTGAACTCCAGTCAC-Index -ATCTCGTATGCCGTCTTCTGCTTG.

5'P indicates the presence of a 5'phosphate group.

Universal adapter:

AATGATACGGCGACCACCGAGATCTACACTCTTTCCCTACACGACGCTCTTCCGATC*T.

* indicates the presence of a phosphorothioate bond between the last C and T.

*Library amplification:* 19.5 µl of sample were mixed with 1µl of a 1:10 dilution of Quant-iT Picogreen dye (Invitrogen, Cat# P11496), 25µl of KAPA HiFi HotStart 2× master Mix (Kapa Bioscience Cat# KK2502), 1µl of 10µM PCR 1.0 and 1µl of PCR 2.0 primers (See below). The amplification mix was distributed in two wells of a LightCycler 384 plate (Roche, Cat# 4729749001) and on a LightCycler 480 II instrument (Roche, Cat# 05015243001) using the following program: 1' at 98°C; N cycles: 10" at 98°C; 20" at 64°C; 45" at 72°C. The number of cycles N was determined on real time by monitoring the fluorescence such that the amplification was stopped during the exponential phase. Samples were removed from the plate and purified with 70 µl SPRI beads without isopropanol, and eluting in 40µl of water. 1µl was used to measure the DNA concentration with a Qubit 3 and the provided reagents (Invitrogen, Cat#Q33218). 1 ng of DNA was used to check fragment size with a D1000 High Sensitivity Screentape and appropriate reagents (Agilent, Cat# 5067-5584, Cat# 5067-5585) on an Agilent 2200 Tapestation.

*Library sequencing:* the libraries were sequenced (paired-end 150bp reads) by Novogene Co. Ltd.

**5/ Gene expression analysis by RNA-seq.**

2 x $10^5$ EKOiE or EKOiE NrKO ES cells were plated in individual wells of 6-well plates and cultured in the presence or absence of 1µg/ml doxycycline (Sigma, Cat# I5148) for 2 days before RNA extraction with 500ul TRIzol (ThermoFisher, Cat# 15596026), according to the manufacturer's protocol. To eliminate any genomic DNA contamination, this was followed by an additional DNAse I treatment (Qiagen, Cat#

79254) for 20min at 37°C followed by phenol:chloroform purification. RNAs were resuspended in Ultrapure DNAse/RNase Free Distilled Water (Thermo, Cat# 10977035). Stranded, poly-A selected RNA-seq libraries were prepared and sequenced (paired-end 150bp reads) by Novogene Co Ltd.

**6/ Western Blot.**

For Western Blot analysis cell pellets corresponding to $10^6$ cells were resuspended in 100 µl RIPA Buffer (Tris/Cl pH 7.5 10mM, NaCl 150mM, EDTA 0.5mM, SDS 0.1%, Triton X-100 1%, Deoxycholate 1%) supplemented with 1× protease inhibitor cocktail (PIC-Roche, Cat# 04 693 116 001) and incubated for 2h in the presence of 2500 U/ml benzonase (Sigma, Cat# E1014). After centrifugation (10 min, full speed, 4 °C), the supernatant was recovered and mixed with an equal volume of 2x Laemmli Sample Buffer (BIO-RAD, Cat# 161-0737) containing β-mercaptoethanol and boiled for 10 min at 95°C and centrifuged for 10 min at maximum speed at room temperature. 20 µl per sample was loaded on 10% Mini-PROTEAN® TGX Stain-Free™ Protein Gels, 10 well, 30 µl (BIO-RAD, Cat# 4568033) and run in 1× SDS-Running Buffer (250 mM Tris/ 1.92 M Glycine/1% SDS) at 10-20 mA using the Mini-PROTEAN Tetra System (BIO-RAD). Proteins were transferred on a Protran nitrocellulose membrane (Amersham, Cat# 10600003) for 1 hour at 300mA using the wet transfer system (BIO-RAD) in 1× Transfer Buffer (10× 0.25M Tris/ 1.92M Glycine) prepared with a final concentration of 20% Ethanol. Membranes were blocked in PBST (PBS 0.1% Tween-20) 5% BSA for 1 hour at room temperature and incubated over night at 4°C with primary antibodies (diluted in PBST 5% BSA). Excess antibodies were washed with PBST (5 washes, 5 min each) and incubated for 1 hour at room temperature in secondary antibodies Alexa Fluor dye or HRP-conjugated (diluted in PBST 5% BSA). Membranes were washed 5 times, 10 min each at room temperature and, for HRP antibodies, incubated with PIERCE ECL2 Western Blotting Substrate (Thermo Scientific, Cat# 80196) 5 min in dark. After excess reagent was removed, proteins were visualised using the BIO-RAD Chemidoc MP Imaging System and processed using the Image Lab Software (BIO-RAD).
Antibodies used:
Primaries: Rabbit polyclonal anti-Lamin B1 (Abcam, Cat# ab16048) (1:10000), goat polyclonal anti-Sox2 (R&D, Cat# AF2018) (1:1000), anti-Esrrb mouse monoclonal (Perseus Proteomics, Cat# H6-705-00) (1:1000), anti-Nanog rabbit polyclonal (Cosmobio, Cat# REC-RCAB001P) (1:1000); anti-Oct4 rabbit polyclonal (Abcam Cat # ab19857) (1:1000). Secondaries: anti-Rabbit IgG-HRP (Thermo Fisher, Cat# RB230254) (1:5.000), Alexa Fluor 488 AffiniPure Donkey Anti-Mouse IgG (H+L) (Jackson ImmunoResearch, Cat #715-545-150); Alexa Fluor 647 AffiniPure Donkey Anti-Goat IgG (H+L) (Jackson ImmunoResearch, Cat # 705-605-147).

**7/ Computational Methods.**

***7a/ Data and availability.***
A total of 69 ChIP-seq and 20 RNA-seq libraries were generated and sequenced: ChIP-seq was performed in triplicates; FCS/L RNA-seq in duplicates; 2i/L RNA-seq in triplicates. All these datasets are available upon request (will be available in GEO after peer-review publication).

***7b/ ChIP-seq analyses.***
Paired end reads were trimmed by aligning read pairs to discover regions of reverse complementarity surrounded by adapters, alignment and trimming were performed with the BioSequences package for Julia 0.6 (Bezanson et al., 2017). Reads were aligned with Bowtie2 (Langmead and Salzberg, 2012) to the mm10 genome, with options "-k 10". Reads were additionally filtered for those with a single discovered alignment (in Bowtie2 "k" mode this is mapping quality = 255) and an edit distance less than 4. Duplicate reads (those aligning with same left-right coordinate) were collapsed into one. Peaks were called against relevant inputs for all samples using MACS2 (Feng et al., 2012) with "callpeak -q 0.2 -g mm". Peaks intersecting with the mm10 blacklist (Encode_Project_Consortium, 2012) were

excluded. We further excluded an outlying replicate for Sox2 in 2i/LIF -E-N identified by principal components analysis and the fraction of reads in peaks of 5.3% as compared to 17.9% and 21.9% for the other two replicates. To determine a set of candidate binding regions for each TF we required that a peak must be called in all replicates of a given condition. We then merged the peaks of each TF analysed in FCS/L and 2i/L, respectively, to obtain regions were multiple TF bind. To quantify ChIP signal at each of these merged regions we took the mean signal over replicates from the original TF peak if present, and took the mean signal over the merged interval if a TF peak was not present. Read counts for boxplots correspond to the midpoint of the paired-end fragment normalised to units of reads per million (RPM). For k-means clustering, as offered by the Clustering package of Julia (Bezanson et al., 2015), we selected regions in which a peak was called for all four TF (Esrrb, Oct4, Sox2 and Nanog) and normalized binding levels at each region for each factor to the condition displaying maximal binding. To determine the cluster number we employed the proximity enrichment to differentially expressed genes (DEGs) described below, and found k=5 was the first k at which the presented ChIP region classes were visible and the enrichment of DEGs in proximity to ChIP-seq peaks robust. The clusters were further complemented by differential binding analysis using DESeq 2 (Love et al., 2014). We set size factors according to total mapped library reads, and employed design `~ChipInput + Esrrb + Nr5a2 + Esrrb:Nr5a2` aimed at determining the effect of Esrrb and Nr5a2 depletion and their interaction over input. We tested for Esrrb loss, Nr5a2 loss and the loss of both against the presence of both TFs. Gene Ontology analyses of each cluster were made with GREAT using standard parameters. De-novo motif discovery was performed on the ensemble of Esrrb/Nr5a2 bound regions in FCS/L using the Regulatory Sequence Analysis Tools (RSAT) through their web-based interface with standard parameters (rsat.sb-roscoff.fr/)(Nguyen et al., 2018) . A matrix for the highest ranking discovered motif was exported, trimmed and plotted using the TFBSTools R package (Tan and Lenhard, 2016). Two PFM matrixes were created to reflect a perfect match to the consensus sequence TCA AGG TCA or TCA AGG CCA, and occurrence of these motifs in Esrrb and Nr5a2 bound regions was determined using TFBSTools. Only matches displaying 0 or 1 mismatch to the consensus, while requiring an exact match to either variant of the 7$^{th}$ base, were considered, and the highest scoring matches to the motif was retained for each region. ChIP-seq data visualization was made using the following R packages and assisted by ggplot2 (Wickham, 2016): ChIP-seq profiles were extracted from Bigwig files using rtracklayer (Lawrence et al., 2009), smoothed with zoo (Zeileis and Grothendieck, 2005) and plotted using the Gviz (Hahne, 2016) and GenomicFeatures (Lawrence et al., 2013) R packages; enrichment heatmaps and metaplots were computed with bamsignals (Mammana, 2020); Venn diagrams were made with eulerr (Larsson, 2020), heatmaps were made with ComplexHeatmap (Gu et al., 2016b).

***7c/ RNA-seq analyses.***

Stranded paired end RNA-seq reads were aligned to the mm10 genome using STAR (Dobin et al. 2013) and quantified by RSEM (Li and Dewey, 2011) using the RSEM-STAR pipeline, with additional options "--seed 1618 --calc-pme --calc-ci --estimate-rspd --paired-end". RSEM estimated read counts per sample were rounded for use with DESeq2 (Love et al., 2014). Genes with at least 20 raw counts in all replicates of at least one condition were considered for differential expression analysis. For all differential expression tests DESeq2 was run without independent filtering; genes considered with absolute FC > 1.5 and FDR < 0.01 were considered as differentially expressed.. Gene Ontology analyses were done in geneontology.org (PANTHER) with standard parameters. To determine enrichments of each group of differentially expressed genes in proximity to the ChIP-seq clusters, we calculated Hypergeometic right tail p-values for for the association between differentially expressed genes within xbp of a ChIP-seq peak of a cluster to a background of all genes clustered within xbp of a cluster peak, for x in [1, 1e+6] bp, using the Julia package ProximityEnrichment.jl (https://github.com/owensnick/ProximityEnrichment.jl). Data visualization was made in R using ggplot2(Wickham, 2016) and ComplexHeatmaps packages (Gu et al., 2016b).

***7d/ Comparisons to published datasets.***

Oct4 ChIP-seq datasets from (Buecker et al., 2014) and (Festuccia et al., 2018) were obtained through the European Nucleotide Archive database and aligned with Bowtie2 (Langmead and Salzberg, 2012) to the mm10 genome, with default options. Output sam files were converted to bam, sorted, and indexed using Samtools (Li et al., 2009). Coverage in the set of Esrrb dependent or independent regulatory regions identified in this study was quantified in each external dataset using the R packages bamsignals (Mammana, 2020), Rsamtools (Morgan, 2020), and GenomicRanges (Lawrence et al., 2009), and data plotted with ggplot2 (Wickham, 2016).

Processed data for RNA-seq correlations were obtained from supplementary tables available in the following publications, and differentially genes selected according to the following criteria:

- (Dunn et al., 2019):
  Up in 2i/LIF vs PD/LIF (Activated by CH) – Fold change > 2, Absolute value in 2i/LIF> 5 FPKM (after averaging replicates)
  Down in 2i/LIFL vs PD/LIF (Repressed by CH) – Fold change < 0.5, Absolute value in PD/LIF > 5 FPKM (after averaging replicates)
- (Ye et al., 2013):
  Up in LIF+CH / LIF (Activated by CH) – Fold change >2, detection p-value <0.1
  Down in LIF+CH / LIF (Repressed by CH) – Fold change <0.5, detection p-value <0.1
- (Martello et al., 2013):
  Upregulated after LIF treatment (1h) in Stat3 wild-type cells cultured in 2i – p-value < 0.05
  Downregulated after LIF treatment (1h) in Stat3 wild-type cells cultured in 2i – p-value < 0.05
- (Yi et al., 2011):
  Upregulated in $Tcf3^{-/-}/Tcf3^{+/+}$ – Fold change <0.5 (after averaging replicates)
  Downregulated in $Tcf3^{-/-}/Tcf3^{+/+}$ – Fold change >1.5 (after averaging replicates)
- (Boroviak et al., 2015):
  Upregulated in ICM3.5/Epi5.5 – Fold change > 2, Absolute value > 5 (after averaging replicates)
  Downregulated in ICM3.5/Epi5.5 – Fold change < 0.5, Absolute value > 5 (after averaging replicates)
  Upregulated in Epi4.5/Epi5.5 – Fold change > 2, Absolute value > 5 (after averaging replicates)
  Downregulated in Epi4.5/Epi5.5 – Fold change < 0.5, Absolute value > 5 (after averaging replicates)
- (Buecker et al., 2014):
  Upregulated in ESC/EpiLC – Fold change > 2, Absolute value > 5 FPKM
  Downregulated in ESC/EpiLC – Fold change < 0.5, Absolute value > 5 FPKM
- (Festuccia et al., 2018):
  Upregulated in $Esrrb^{High}/Esrrb^{Negative}$ – Fold change > 2 (after averaging replicates)
  Downregulated in $Esrrb^{High}/Esrrb^{Negative}$ – Fold change < 0.5 (after averaging replicates)

Pearson correlation coefficients were calculated based on the fold changes in expression of each gene in the external datasets and the fold change in expression observed after depletion of Esrrb, Nr5a2 or both Esrrb and Nr5a2 in FCS/LIF or 2i/LIF. Heatmaps displaying the correlation between datasets were generated using the ComplexHeatmap R package(Gu et al., 2016b).

Processed data for RNA-seq analysis of Esrrb and Nr5a2 relative levels of expression was obtained from supplementary tables available in the following publications: (Atlasi et al., 2019; Buecker et al., 2014; Cruz-Molina et al., 2017; Dunn et al., 2019; Ficz et al., 2013; Gu et al., 2016a; Hsu et al., 2019; von Meyenn et al., 2016). Processed single cell RNA-seq (Kolodziejczyk et al., 2015) was obtained from the ESpresso database (https://espresso.teichlab.sanger.ac.uk/), and data imported in and plotted with the FlowJo software suite.

**Supplementary references**


- Atlasi, Y., Megchelenbrink, W., Peng, T., Habibi, E., Joshi, O., Wang, S.Y., Wang, C., Logie, C., Poser, I., Marks, H., et al. (2019). Epigenetic modulation of a hardwired 3D chromatin landscape in two naive states of pluripotency. Nat Cell Biol 21, 568-578.
- Bezanson, J., Edelman, A., Karpinski, S., and Shah, V. (2015). Julia: A fresh approach to numerical computing. https://arxivorg/abs/14111607.
- Boroviak, T., Loos, R., Lombard, P., Okahara, J., Behr, R., Sasaki, E., Nichols, J., Smith, A., and Bertone, P. (2015). Lineage-Specific Profiling Delineates the Emergence and Progression of Naive Pluripotency in Mammalian Embryogenesis. Dev Cell 35, 366-382.
- Buecker, C., Srinivasan, R., Wu, Z., Calo, E., Acampora, D., Faial, T., Simeone, A., Tan, M., Swigut, T., and Wysocka, J. (2014). Reorganization of enhancer patterns in transition from naive to primed pluripotency. Cell Stem Cell 14, 838-853.
- Cruz-Molina, S., Respuela, P., Tebartz, C., Kolovos, P., Nikolic, M., Fueyo, R., van Ijcken, W.F.J., Grosveld, F., Frommolt, P., Bazzi, H., et al. (2017). PRC2 Facilitates the Regulatory Topology Required for Poised Enhancer Function during Pluripotent Stem Cell Differentiation. Cell Stem Cell 20, 689-705 e689.
- Dunn, S.J., Li, M.A., Carbognin, E., Smith, A., and Martello, G. (2019). A common molecular logic determines embryonic stem cell self-renewal and reprogramming. EMBO J 38.
- Encode_Project_Consortium (2012). An integrated encyclopedia of DNA elements in the human genome. Nature 489, 57-74.
- Feng, J., Liu, T., Qin, B., Zhang, Y., and Liu, X.S. (2012). Identifying ChIP-seq enrichment using MACS. Nat Protoc 7, 1728-1740.
- Festuccia, N., Halbritter, F., Corsinotti, A., Gagliardi, A., Colby, D., Tomlinson, S.R., and Chambers, I. (2018). Esrrb extinction triggers dismantling of naive pluripotency and marks commitment to differentiation. EMBO J 37.
- Ficz, G., Hore, T.A., Santos, F., Lee, H.J., Dean, W., Arand, J., Krueger, F., Oxley, D., Paul, Y.L., Walter, J., et al. (2013). FGF signaling inhibition in ESCs drives rapid genome-wide demethylation to the epigenetic ground state of pluripotency. Cell Stem Cell 13, 351-359.
- Gu, K.L., Zhang, Q., Yan, Y., Li, T.T., Duan, F.F., Hao, J., Wang, X.W., Shi, M., Wu, D.R., Guo, W.T., et al. (2016a). Pluripotency-associated miR-290/302 family of microRNAs promote the dismantling of naive pluripotency. Cell Res 26, 350-366.
- Gu, Z., Eils, R., and Schlesner, M. (2016b). Complex heatmaps reveal patterns and correlations in multidimensional genomic data. Bioinformatics 32, 2847-2849.
- Hahne, F. (2016). Visualizing Genomic Data Using Gviz and Bioconductor, in Statistical Genomics. Methods and Protocols. Anticancer Res 36, 3224.
- Hsu, J., Arand, J., Chaikovsky, A., Mooney, N.A., Demeter, J., Brison, C.M., Oliverio, R., Vogel, H., Rubin, S.M., Jackson, P.K., et al. (2019). E2F4 regulates transcriptional activation in mouse embryonic stem cells independently of the RB family. Nature Communications 10.
- Kolodziejczyk, A.A., Kim, J.K., Tsang, J.C., Ilicic, T., Henriksson, J., Natarajan, K.N., Tuck, A.C., Gao, X., Buhler, M., Liu, P., et al. (2015). Single Cell RNA-Sequencing of Pluripotent States Unlocks Modular Transcriptional Variation. Cell Stem Cell 17, 471-485.
- Langmead, B., and Salzberg, S.L. (2012). Fast gapped-read alignment with Bowtie 2. Nat Methods 9, 357-359.
- Larsson, J. (2020). eulerr: Area-Proportional Euler and Venn Diagrams with Ellipses. R package version 6.1.0.

- Lawrence, M., Gentleman, R., and Carey, V. (2009). rtracklayer: an R package for interfacing with genome browsers. Bioinformatics 25, 1841-1842.
- Lawrence, M., Huber, W., Pages, H., Aboyoun, P., Carlson, M., Gentleman, R., Morgan, M.T., and Carey, V.J. (2013). Software for computing and annotating genomic ranges. PLoS Comput Biol 9, e1003118.
- Li, B., and Dewey, C.N. (2011). RSEM: accurate transcript quantification from RNA-Seq data with or without a reference genome. BMC Bioinformatics 12, 323.
- Li, H., Handsaker, B., Wysoker, A., Fennell, T., Ruan, J., Homer, N., Marth, G., Abecasis, G., Durbin, R., and Genome Project Data Processing, S. (2009). The Sequence Alignment/Map format and SAMtools. Bioinformatics 25, 2078-2079.
- Love, M.I., Huber, W., and Anders, S. (2014). Moderated estimation of fold change and dispersion for RNA-seq data with DESeq2. Genome Biol 15, 550.
- Mammana, A. (2020). bamsignals: Extract read count signals from bam files. R package version 1.20.0.
- Martello, G., Bertone, P., and Smith, A. (2013). Identification of the missing pluripotency mediator downstream of leukaemia inhibitory factor. EMBO J 32, 2561-2574.
- Morgan, M. (2020). Rsamtools: Binary alignment (BAM), FASTA, variant call (BCF), and tabix file import. R package version 2.4.0.
- Nguyen, N.T.T., Contreras-Moreira, B., Castro-Mondragon, J.A., Santana-Garcia, W., Ossio, R., Robles-Espinoza, C.D., Bahin, M., Collombet, S., Vincens, P., Thieffry, D., et al. (2018). RSAT 2018: regulatory sequence analysis tools 20th anniversary. Nucleic Acids Res 46, W209-W214.
- Tan, G., and Lenhard, B. (2016). TFBSTools: an R/bioconductor package for transcription factor binding site analysis. Bioinformatics 32, 1555-1556.
- von Meyenn, F., Iurlaro, M., Habibi, E., Liu, N.Q., Salehzadeh-Yazdi, A., Santos, F., Petrini, E., Milagre, I., Yu, M., Xie, Z., et al. (2016). Impairment of DNA Methylation Maintenance Is the Main Cause of Global Demethylation in Naive Embryonic Stem Cells. Mol Cell 62, 983.
- Wickham, H. (2016). ggplot2: Elegant Graphics for Data Analysis. Springer-Verlag New York.
- Ye, S., Li, P., Tong, C., and Ying, Q.L. (2013). Embryonic stem cell self-renewal pathways converge on the transcription factor Tfcp2l1. EMBO J 32, 2548-2560.
- Yi, F., Pereira, L., Hoffman, J.A., Shy, B.R., Yuen, C.M., Liu, D.R., and Merrill, B.J. (2011). Opposing effects of Tcf3 and Tcf1 control Wnt stimulation of embryonic stem cell self-renewal. Nat Cell Biol 13, 762-770.
- Zeileis, A., and Grothendieck, G. (2005). zoo: S3 infrastructure for regular and irregular time series. J Stat Softw 14.
